# Injury-free priming: induction of Primary Afferent Collateral Sprouting in uninjured sensory neurons *in vivo* primes them for enhanced axon outgrowth *in vitro*

**DOI:** 10.1101/463935

**Authors:** Sara Soleman, Jeffrey C. Petruska, Lawrence D.F. Moon

## Abstract

Prior “conditioning” nerve lesions can prime DRG neurons for enhanced axon regeneration. Here, we tested the hypothesis that adult DRG neurons can be primed for axon elongation in vitro without axonal injury by prior induction of Primary Afferent Collateral Sprouting (PACS) in vivo. Thoracic cutaneous nerves (T9, T10, T12, T13 but not T11) were transected to create zones of denervated skin. Neurons from the uninjured T11 DRG underwent PACS within the skin, as demonstrated by the expansion of its zones responsive to pinch up to 14 days. At 7 or 14 days after induction of collateral sprouting, DRG neurons were dissociated and cultured for 18 hours in defined media lacking neurotrophins and growth factors. Neurons from the uninjured T11 DRG had longer mean neurite lengths than neurons from naïve DRG. A larger proportion of neurons from the uninjured T11 DRG showed an elongating or arborizing phenotype than neurons from naïve DRG. Transcriptomic analysis of the uninjured T11 DRG and denervated/reinnervated skin reveal regulation of receptor/ligand systems and regulators of growth during collateral sprouting. For example, the glial cell-derived neurotrophic family ligands Artemin and Persephin were upregulated in denervated skin after 7 and/or 14 days. We suggest that extracellular cues in denervated skin modify the intrinsic growth program of uninjured DRG neurons that enhances their ability to elongate or arborize even after explantation. Collectively, these data confirm that induction of collateral sprouting does not induce an injury response yet primes many of these uninjured neurons for in vitro axon growth.

## 1. Introduction

After injury, axons in the central nervous system (CNS) extend collateral branches and sprout considerably but do not elongate long distances back to their original targets (Rosenzweig et al,. 2010). In contrast, after injury to the peripheral branch of the dorsal root ganglia (DRG), many sensory axons eventually elongate axons back to their original targets although this does not occur after injury to the central branch of the DRG (Snider et al., 2002). Axons from sensory neurons or motor neurons can be primed to elongate more effectively when a conditioning stimulus is given prior to an injury (Gutmann, 1942). For example, a conditioning injury to the peripheral branch of a DRG increases the number of regenerating sprouts and enhances their elongation (McQuarrie et al., 1977; McQuarrie, 1985; Richardson and Verge, 1987; Jenq et al., 1988). Indeed, a conditioning lesion has been shown to prime the cell body of axotomized neurons by activating an intrinsic transcriptional program that switches the neuron from an arborizing state into an elongating state (Smith and Skene, 1997; Qiu et al 2005): for example, after explantation, dissociated neurons from DRG of unoperated rats arborize after a delay whereas deliberately pre-injured DRG neurons initiate axon growth more rapidly in an elongating mode rather than an arborizing mode (Smith and Skene, 1997).

Here, we set out to test the hypothesis that DRG neurons can be primed to undergo elongation without prior injury. We took advantage of an existing, well characterised model for studying Primary Afferent Collateral Sprouting of <u>uninjured</u> sensory axons (Devor et al., 1979; Diamond et al., 1976; Bisby et al., 1996; Petruska et al., 2014). Cutaneous axons from one thoracic DRG almost exclusively project to a single dermatome (Ygge, 1984). One can induce sprouting of uninjured axons from a thoracic DRG into adjacent dermatomes by transecting neighbouring thoracic cutaneous nerves (Figure 1); very few, if any, neurons in the spared DRG are injured (Devor et al., 1979; Diamond et al., 1976; Bisby et al., 1996). Over a few weeks, these uninjured neurons sprout axons into adjacent, denervated dermatomes (Diamond et al., 1992b; Doucette and Diamond, 1987; Nixon et al., 1984).

**Figure 1.**
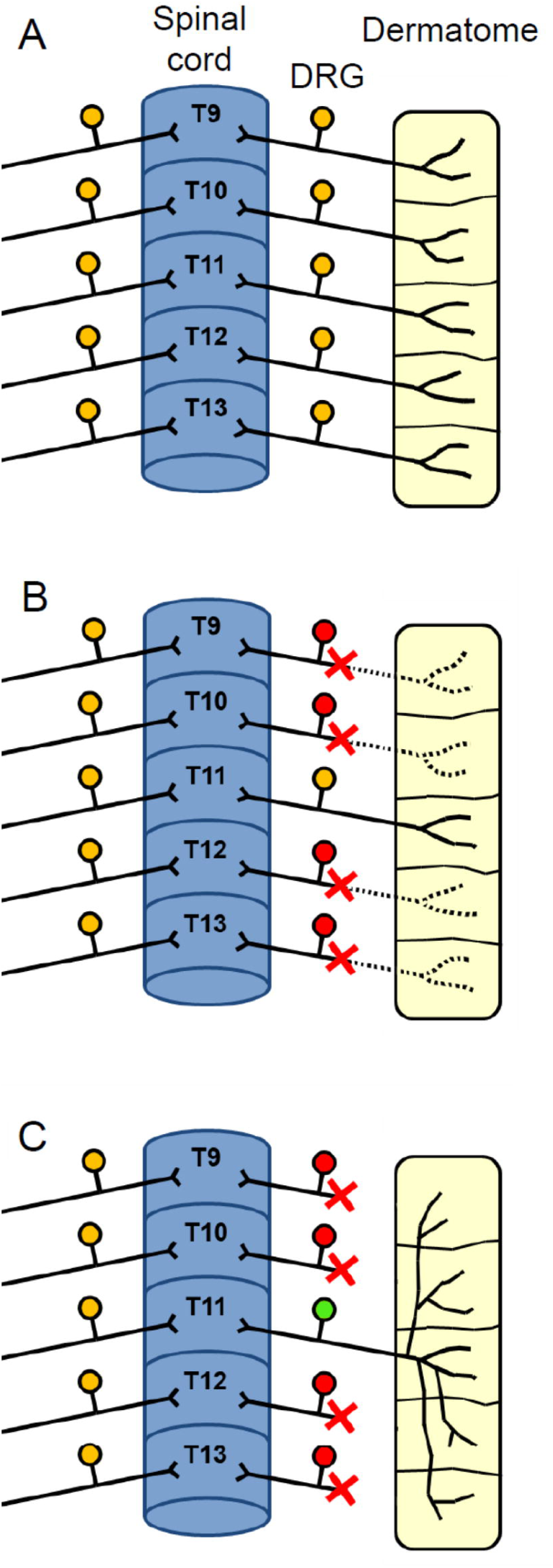
Schematic showing the central and peripheral branches of thoracic DRG neurons before and after induction of PACS. **(A)** Thoracic DRG neurons (orange) exhibit a 1:1:1 correspondence between a spinal cord segment (blue), DRG and the dermatome they innervate (yellow) without involvement in a nerve plexus. **(B)** Transection and ligation of cutaneous nerves (red crosses) neighbouring the spared T11 nerve (orange) leaves a cutaneous sensory field innervated by the uninjured DRG neurons and surrounded by a vast area of denervation. **(C)** Over time, the peripheral branch of the spared, uninjured “T11 PACS DRG” (green) undergoes primary afferent collateral sprouting to innervate previously denervated areas to restore sensorimotor reflexes and function. In this study the “contralateral T11 DRG” was used as a control in some cases. The “contralateral-to-transection DRGs” (T9, T10, T12, T13) were pooled and also used as controls.

Accordingly, we induced PACS *in vivo* in uninjured T11 DRG neurons. Seven or fourteen days later, uninjured T11 DRG neurons were dissociated and cultured in the absence of neurotrophic factors. We compared these uninjured T11 DRG neurons with neighbouring DRG with nerve injury and with neurons from T11 DRG of unoperated rats. We examined the question of whether neurons can be primed for axonal elongation without nerve injury.

## 2. Methods

### Surgery

All procedures were in accordance with the UK Home Office and Animals (Scientific Procedures) Act of 1986, associated guidelines and the EU Directive 2010/63/EU for animal experiments. Ethical approval was given by the King’s College London Animal Welfare and Ethical Review Board and work was carried out under a Project Licence from the UK Home Office (70/6851) and by Personal Licences held by Dr Lawrence Moon and Sara Soleman.

Adult female Sprague-Dawley rats (260-310 g, *n*=7, Harlan, UK) were anesthetized with isoflurane (3% in O_2_ for induction; 1.5-2.0% for maintenance). Fur was removed from the thoracolumbar region of the back and a 4 to 5 cm longitudinal incision of the skin was made approximately 1 cm to the right of the midline and clipped back. The off-midline incision was performed to avoid cutting left-side sensory axons which induces an injured state (Hill et al, 2010; Rau et al., 2016). Blunt dissection of subcutaneous fascia exposed the array of dorsal cutaneous nerves and lateral cutaneous nerves. Thoracic (T) T9, T10, T12 and T13 dorsal and lateral cutaneous nerves were identified (Figure 1A) and carefully freed from both connective tissue and the latissimus dorsi muscle wall. These cutaneous nerves were then transected a few millimetres from the body wall and ligated with 7-0 Ethilon sutures (Figure 1B). This procedure successfully disallows nerve regeneration to the skin in almost every instance (Diamond et al., 1992b). The T11 dorsal and lateral cutaneous nerves were left intact, with the T11 sensory field “isolated” within a relatively vast area of denervated skin (Figure 1B) creating a “spared dermatome”. The dorsal and lateral cutaneous nerves were both transected in this experiment in order to provide a greater area of neighbouring denervation around the spared T11 dermatome than if only the dorsal cutaneous nerves were transected. The skin incision was then closed. Before animals were allowed to recover from anaesthesia, mechanonociceptive fields were mapped as described below. Rats were terminally anesthetized at day 7 (n=4) or day 14 (n=3) because it is known that PACS occurs during this time frame. Analgesic (buprenorphine, 0.01 mg/kg subcutaneously) was given for 3 days after surgery.

At the end of the survival period (7 or 14 days post-surgery), animals were deeply anesthetized and the DRGs were retrieved either unfixed or fixed (see below). To ensure isolation of the correct DRGs, the spared and transected cutaneous nerves were traced back through the *latissimus dorsi* muscle to the parent ganglion via microdissection.

This surgery provided both uninjured neurons undergoing PACS (T11 DRG) and injured neurons (T9, 10, 12, 13 DRG). We retrieved the corresponding contralateral DRGs to use as a comparison group. However, we must note that these contralateral DRGs likely contain a small number of neurons that have been injured by the incision (Hill et al 2010; Rau et al 2016) and some innervating the peri-incisional skin that may have been induced into the PACS-state by the local inflammation. “Naive” T11 DRG were also retrieved from unoperated rats (n=3).

DRG neurons explanted from naive (unoperated) rats will be referred to as “naive DRG” and “naive neurons”, although clearly the explant procedure induces an injury. T11 “PACS DRG” will be used to refer to uninjured, T11 DRG which underwent PACS prior to explantation. “Transected DRG” will be used to refer to thoracic DRG whose nerves had been transected prior to explantation.

### Behavioural mapping of nociceptive fields

A simple behavioural test involving the *cutaneous trunci* muscle (CTM) reflex system was used to monitor progress of collateral sprouting, synapse formation and functional recovery (Diamond et al., 1987; Doucette and Diamond, 1987; Diamond et al., 1992b). Mechanonociceptive fields were “pinch mapped” under isoflurane anaesthesia, <u>adjusted per rat to a light enough level</u> that enabled brisk but localised reflex responses to pinching of the skin or foot. Fine-toothed forceps were used to pinch the skin which evokes a reflex contraction of the underlying CTM, causing a visible contraction of the skin (Figure 2A, B) (Therieault and Diamond 1998; Petruska et al., 2014; Harrison et al., 2015; Rau et al 2016). The borders between sensitive and insensitive areas of the skin were determined by the respective presence or absence of the CTM response to successive pinch stimuli at approximately 2 to 3 mm intervals. The skin was marked using a permanent marker between unresponsive and responsive test sites, to enable assessment of the field expansion over time (Figure 1C and Figure 2C, D). It should be noted that nociceptive dermatomes overlap such that denervation of at least two adjacent dermatomes was required to generate an adequate “nociceptor-denervated” area of skin. The expansion of the functional CTM reflex receptive field corresponds with the expansion of anatomical innervation fields of the spared/sprouting axons (e.g., Pertens et al., 1999) Diamond et al 1992b). Accordingly, here, primary afferent collateral sprouting was confirmed by “pinch mapping” in all of the rats included in this study (whether for cell culture or for microarray analysis). We have previously quantified the rate of reduction of the denervated zone using other rats (Harrison et al 2015) and these open access data are reproduced here for convenience (Figure 2E).

**Figure 2.**
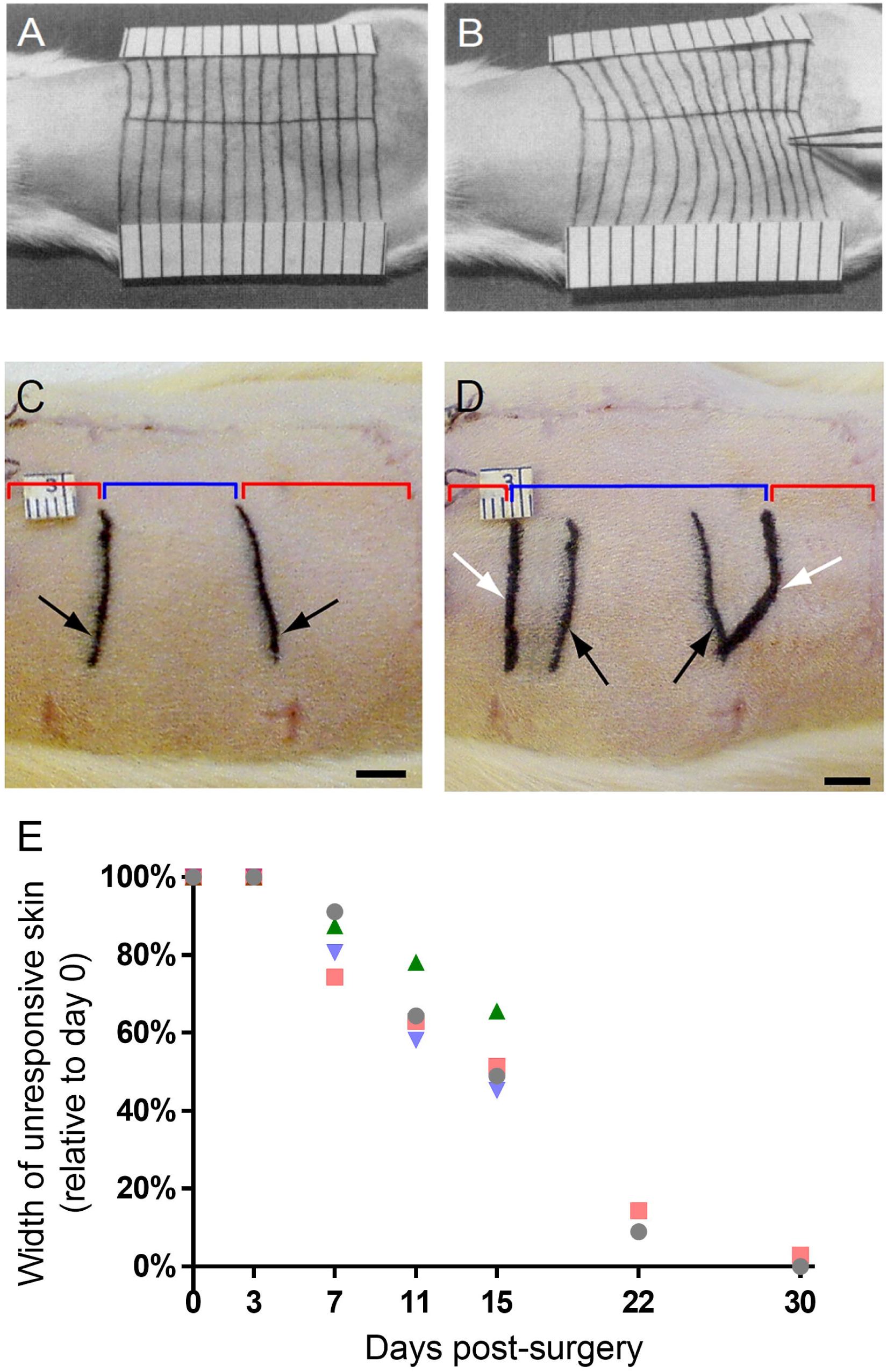
The response to pinching is lost in a denervated zone created by transection of cutaneous nerves but is restored after collateral sprouting by uninjured neurons from the spared T11 DRG. (A) Noxious mechanical stimulation activates the *Cutaneous Trunci* Muscle (CTM) reflex response in a somatotopic manner. (B) Pinch evokes contraction of the skin in normal, lightly anesthetised rats. Images in A and B are from Theriault and Diamond (1988). (C) Following cutaneous nerve transection, behavioural “pinch mapping” reveals an area (outlined in black on the skin) within which the CTM reflex response can be evoked. The blue line and black arrows indicate the responsive “isolated” mechanonociceptive field and red lines denote unresponsive mechanonociceptive fields. (D) At 14 days after surgery, “pinch mapping” in the same rat showed expansion of the zone exhibiting a CTM reflex response (denoted by the blue line and white arrows, with original responsive zone denoted by the black arrows. Regions of denervation had shrunk (denoted by red lines). (C-D) Scale bars: 5 mm. (E) Graph showing the reduction over time in the area of skin unresponsive to pinch, expressed as a percentage of the area of skin that was unresponsive on day 0; n=4 rats; adapted from Harrison et al 2015 (https://creativecommons.org/licenses/by-nc-nd/4.0/).

### Immunolabelling for Activating Transcription Factor 3 to assess injury status of DRG neurons

An additional set of adult female Sprague-Dawley rats (260-310 g) was anesthetized with pentobarbital (65 mg/kg, i.p.). Of these, two rats served as shams and were not operated upon further. In other rats (n=5), sprouting of T11 DRG neurons was induced with surgical isolation of the T11 cutaneous sensory fields by unilateral transection of T9, T10, T12, and T13 of dorsal and lateral cutaneous nerves (Figure 1). In the operated rats, denervation of the dermatomes was confirmed by lack of CTM reflex in response to pinch. We also used CTM reflex testing to confirm success of PACS. One additional rat received transection of the sciatic nerve at mid-thigh level. Analgesic (buprenorphine, 0.01 mg/kg subcutaneously) was given for 3 days after surgery. Seven or Fourteen days after spared-dermatome surgery, or 7 days after sciatic nerve injury, rats were deeply anesthetized with pentobarbital (65 mg/kg, i.p.) and perfused transcardially with saline followed by 4% paraformaldehyde in 0.1 M phosphate buffer. Spinal cord and DRGs were removed and post-fixed in 4% paraformaldehyde overnight. Tissue was transferred to a solution of 20% sucrose (Sigma, Poole, UK) for at least 24 hours at 4°C.

DRG sections were cut at a thickness of 10 μm on a cryostat. Sections of DRG were mounted onto Superfrost Plus slides (BDH, Poole, UK). Sections were air dried and incubated in 2% normal goat serum in 0.1 M Tris-buffered saline (TBS) containing 0.2% Triton X-100 for 60 min. Sections were then incubated in rabbit polyclonal anti-ATF3 antibody (1:500, overnight; Santa Cruz, CA, #C-19, sc-188) in 0.1 M Tris-buffered saline (TBS) containing 0.2% Triton X-100. Sections were washed in TBS containing 0.2% Triton X-100 three times and incubated in Alexa 488-conjugated goat anti-rabbit secondary antibody (1:500, 4 hours; Invitrogen, A-11034) containing bisbenzimide (1 μg/ml). They were washed in TBS containing 0.2% Triton X-100 three times and washed in distilled water. Sections were mounted onto coverslip in Vectashield medium (Vector). Control sections were treated in the same manner, but with 10% normal goat serum replacing the primary antibody. The number of ATF3 positive nuclei per section were counted and averaged across sections per DRG. We analysed a total of four naive T11 DRGs, five T11 PACS DRGs and four DRGs with transected nerves.

### Primary DRG cultures

DRG from T9 to T13 on both the ipsilateral and contralateral side to surgery were collected and placed in individual wells of a 24 well plate containing Hams F-12 media (Life Technologies, Gaithersburg, MD). Ganglia were manually and gently cleared of connective tissue. They were then individually incubated in 0.125% collagenase and 50 µg/ml DNase (Sigma) for 2.5 hr at 37°C in 5% CO_2_ to chemically dissociate and prevent cell clumping during tissue disaggregation. Ganglia were then mechanically dissociated individually by gentle trituration in 1 ml of F-12 media then centrifuged through a 15% BSA cushion at 700 rpm (77 x g) for 7 min. Dissociated neurons were re-suspended in 100 µl of modified Bottenstein and Satos medium (BS): Neurobasal media (Invitrogen) supplemented with 100 units/ml BSA (30%; Sigma), 100 units/ml N2-supplement (Invitrogen), 100 units/ml penicillin and 100µg/ml streptomycin (Invitrogen). This defined medium lacks neurotrophins and growth factors including NGF, BDNF and GDNF. Cells were then plated onto Lab-Tek (Nunc, Fisher Scientific, Loughborough, UK) chamber slides coated with poly-L-lysine (2 mg/ml; Sigma). The dissociation of individual ganglia generally retrieved between 800-1000 cells and the suspension was divided into two chambers to provide low-density cultures. Cultures were incubated for 18 hr at 37°C in a humidified atmosphere containing 5% CO_2_.

### Immunocytochemistry

Cells were fixed with 4% PFA for 20 min at room temperature, then permeabilized with methanol (100%) for 3 min and washed 3 times in PBS. To assess neurite outgrowth, all cells were incubated in the following: mouse anti-βIII tubulin (1:1000; 2 hr; Promega) and goat anti-mouse Alexa 546 (1:1000; 45 min; Molecular Probes, Invitrogen). Lab-Tek chambers were removed and slides were coverslipped with Vectashield mounting medium with DAPI (Vector Labs).

### Quantitative analysis of neurite outgrowth

All images were captured using a Carl Zeiss AxioImager Z1 microscope and analysed by an investigator blinded to different groups. Analysis was organised into the following groups: 1) Spared T11 DRG, 2) Transected DRGs (all values for individual transected DRGs were pooled for analysis), 3) Contralateral spared T11 DRG, and 4) Contralateral-to-transected DRGs (T9, T10, T12, T13 were pooled for analysis).

#### Analysis of neurite outgrowth

The number of sprouting neurons, visualised by βIII tubulin immunoreactivity, were quantified using two parameters: all sprouting neurons with neurites extending longer than 10 µm or greater than 50 µm. The percentage of sprouting neurons was calculated as follows: (number of sprouting neurons/total number of neurons) x 100. The total number of surviving neurons per well ranged from 150 – 500 cells.

#### Sholl analysis

Neurons with projections greater than 50 µm were analysed using Sholl analysis to determine the total neurite length and number of branches at each shell distance. Grayscale 8-bit images were obtained and semi-automatically traced with NIH Image J using the Sholl Analysis plug-in. Analysis parameters for the concentric circles included: starting radius = 50 µm from the centre of the neuron, radius step intervals = 50 µm and ending radius = 900 µm.

Neurons with a mean of <2 branches per 100 µm neurite were categorised as elongating, whereas those with mean >2 branches per 100 µm neurite were categorised as arborizing.

#### Analysis of cell size distributions

Cell size distributions were also determined using the AxioVision LE V4.7.2 (Carl Zeiss) contour tracing software. Cell diameter (µm) was measured in all sprouting neurons extending processes greater than 50 µm and tabulated for each group.

### Real time qRTPCR

In a previous study, to identify genes that were regulated in uninjured T11 DRG neurons during PACS, we performed microarray analysis upon RNA taken from T11 DRG from naïve rats (n=5) and from T11 DRG taken 7 days (n=7) or 14 days (n=7) after induction of PACS (Harrison et al., 2015; Harrison et al., 2016). Microarray analysis was also performed using RNA from those same rats from intact skin (n=6) or 7-day denervated skin (n=5) and 14-day denervated skin (n=5) (Flight et al., 2014). Those datasets may be found at the Gene Expression Omnibus (GEO) with accession numbers GSE72551 (DRG) and GSE54356 (skin).

Here, we used qRTPCR to confirm some of the gene changes identified during microarray analysis. Some results have been presented previously (Harrison et al., 2015). We used antisense RNA (aRNA) left over from the first round of amplification prior to microarray analysis. 200 ng aRNA, 2 μl of Random Primers (250 ng/μl, Invitrogen) and Nuclease-free water (Qiagen) were mixed to total volume 12 μl, and then heated at 70°C for 3 minutes. Then first strand cDNA synthesis was carried out using 1 μl M-MLV Reverse Transcriptase (Ambion Research), 1 μl RNase inhibitor, 2 μl reaction buffer and 4 μl dNTP mix to 20 μl total volume. They were incubated for 1 hour at 42°C, and to inactivate reverse transcriptase heated for 10 minutes at 92°C. To construct standard curve for genes of interest total RNA was extracted from homogenized head of 15/16 days rat embryo (Fischer 344) by TRIzol reagent (Invitrogen). 5 μg total RNA was treated with DNase (Qiagen) and purified (RNeasy Kit, Qiagen). cDNA was produced by using Random Primers (250 ng/μl, Invitrogen) and SuperScript II reverse transcriptase (Invitrogen).

The primer sequences are taken from PrimerBank (Wang and Seed, 2003) or were designed based on the probe set target sequence provided by Affymetrix using Primer3 software (www.genome.wi.mit.edu) (Table 1). Quantitative analysis was carried out by real-time PCR (Rotor-Gene 3000, Corbett Life Science, Sydney, Australia). Each reaction contained cDNA, 0.5 µl of each primer (25 ng/µl), 12.5 µl of 2X SYBR Green PCR Master Mix (AmpliTaq Gold DNA polymerase, reaction buffer, dNTP mix, 2X SYBR Green 1 dye; Applied Biosystems) in 25-μl reaction. Reactions were performed in triplicate for standard curve. GAPDH was selected as control gene because analysis of variance of the microarray data indicated it was invariant in DRG during PACS. Standard curves of genes of interest and endogenous control GAPDH gene were constructed with eight 3-fold dilutions of cDNA from E15/E16 rat head. The amplification program consisted of 1 cycle at 95°C for 10 min followed by 40 cycles of 95°C denaturation for 15 s, 60°C annealing and extension for 60 s. For melting curve analysis the samples were heated from 60°C to 95°C. Single melting curve peaks indicated that a single product was amplified. Amplified products were also run on a 1% agarose gel to show the specificity of the amplicon. After raw data was collected, the data were normalized to account for initial differences in background fluorescence. The fold change was expressed relative to GAPDH value.

**Table 1:**
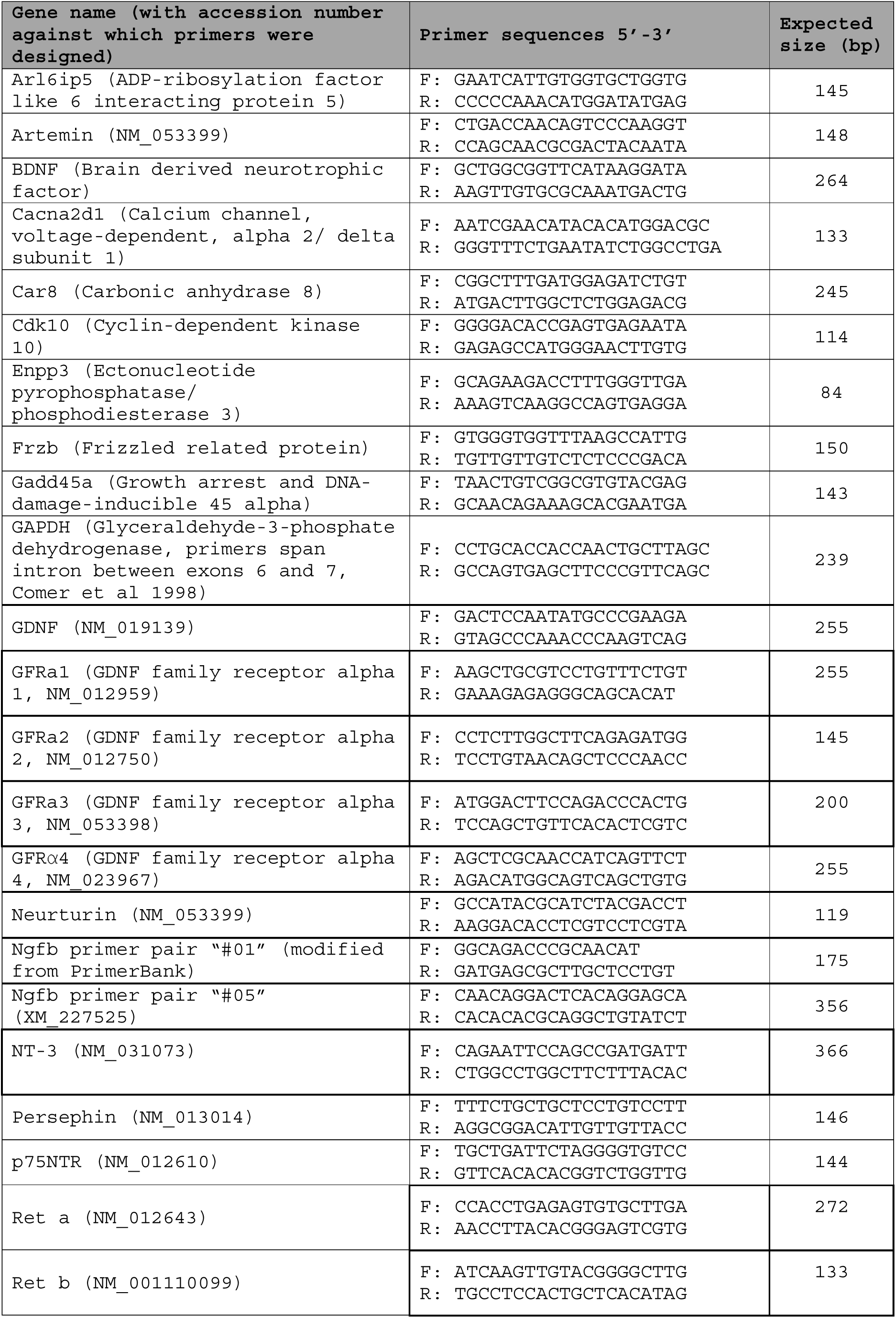
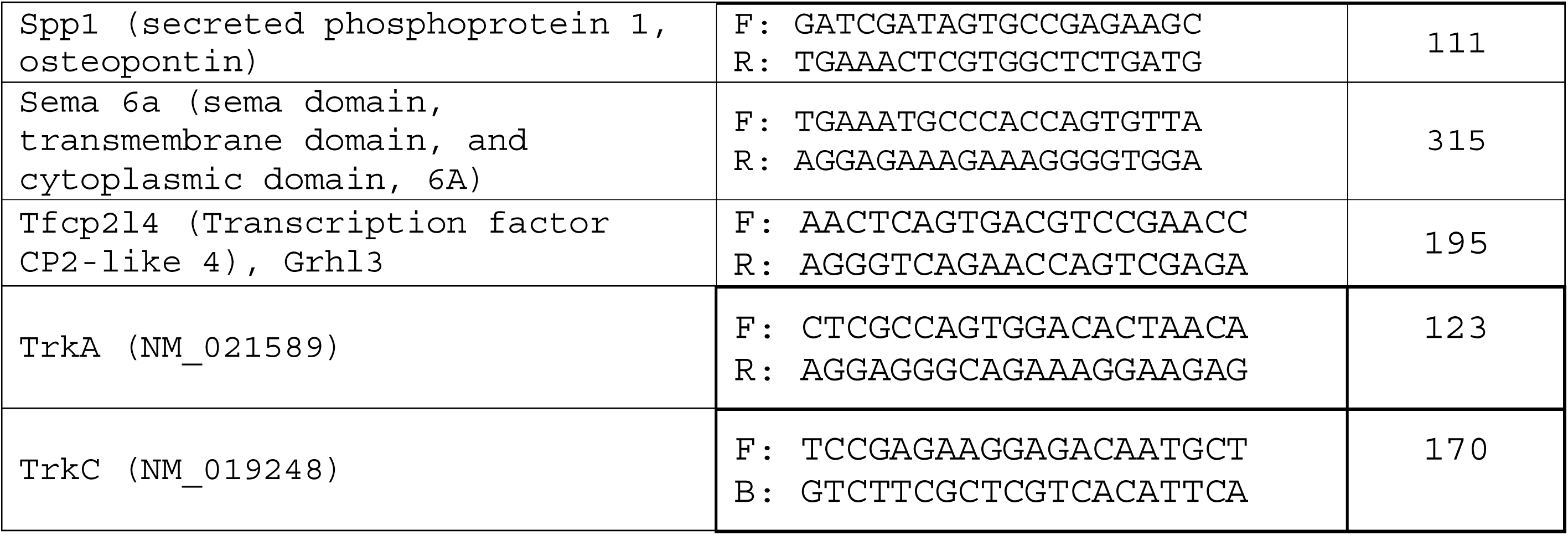
Table shows sequences of primers used during real time RT-PCR. This table lists primers for RTPCR reactions whose results are reported in either this paper or a previous paper (Harrison et al., 2015). Primers shown 5’ to 3’.

### Statistical Analysis

Analysis was performed using SPSS (V17.0). Unless otherwise stated, ‘n’ is the number of independent biological replicates (*i*.*e*., number of rats). The percentage of neurons that sprouted, the percentage of sprouting neurons that were elongating or arborizing, and the percentage of sprouting neurons that were large/small diameter neurons were analysed using either the Mann-Whitney (independent samples from different rats *e*.*g*., naive vs. spared T11 neurons) or Wilcoxon (related samples) tests. Where ANOVAs were significant, pairwise group comparisons were conducted using Fisher’s Least Significant Difference (LSD) *post hoc* tests and exact p-values are reported. To avoid visual and textual clutter, we do not report differences relative to contralateral T11 neurons and to contralateral-to-transected neurons. qRTPCR data were analysed using ANOVA followed by LSD *post hoc* tests. In all cases, *post hoc* tests were reported only when the ANOVA was significant (p<0.05). Scatterplots were generated using GraphPad Prism v8. Data are presented as means ± SEM and asterisks indicate significance as follows: \**p*≤0.05; \*\**p*≤0.01; \*\*\**p*≤0.001.

## Results

### Spared dermatome surgery induced expansion of mechanonociceptive “pinch” fields by uninjured neurons

Mechanonociceptive fields were “pinch mapped” by evoking a reflex in the *cutaneous trunci* muscle (CTM) which results in a visible contraction of the overlying skin (Figure 2A, B). As expected, immediately after surgery, the CTM reflex response was present in the skin innervated by the spared T11 cutaneous nerves (zone shown in blue; Figure 2C) and absent in adjacent zones denervated by transection of cutaneous nerves (zone shown in red; Figure 2C). At 14 days after induction of PACS, behavioural mapping of borders between responsive and unresponsive areas revealed receptive field expansion (Figure 2D; white arrows) relative to previous borders (black arrows). This expansion was consistent with previous studies of Primary Afferent Collateral Sprouting by intact fibers (Diamond et al., 1992b; Doucette and Diamond, 1987; Nixon et al., 1984). In a previous study, we documented the reduction in the area of denervated skin over time (Figure 2E, adapted from Harrison et al 2015).

In that study (Harrison et al., 2015), we obtained microarray data which show that this surgical procedure does not evoke expression of injury-related transcripts in the DRG of the spared dermatome (T11). For example, the level of the injury-related transcript *Activating Transcription Factor 3* (*ATF3*) remained very low and there was no difference in ATF3 mRNA levels between naïve rats and rats at 7 or 14 days after induction of PACS (Figure 3A; ANOVA p>0.05). Very low expression of ATF3 was confirmed in a separate set of DRG by immunolabelling for ATF3. As expected, T11 DRGs from naïve rats contained few or no neurons that were positive for ATF3 (Figure 3C). Very few neurons from T11 DRG taken 7 or 14 days after induction of PACS were positive for ATF3 (Figure 3B, E and F). As positive controls, we immunolabelled sections from adjacent thoracic DRG whose lateral and cutaneous nerves had been cut 7 or 14 days previously and also a lumbar DRG whose sciatic nerve had been transected seven days previously: many neurons showed expression of ATF3 protein, as expected (Figure 3D) (Hill et al., 2010). Importantly, in both the microarray and immunolabelling studies, we used CTM reflex testing to verify expansion of receptive fields in these rats to prove that they had undergone PACS *in vivo* prior to removal of tissues. These data show that T11 DRG neurons from rats undergoing PACS do not express ATF3: this confirms that our surgical procedure does not injure the T11 DRG neurons before they are explanted for culturing (see below).

**Figure 3:**
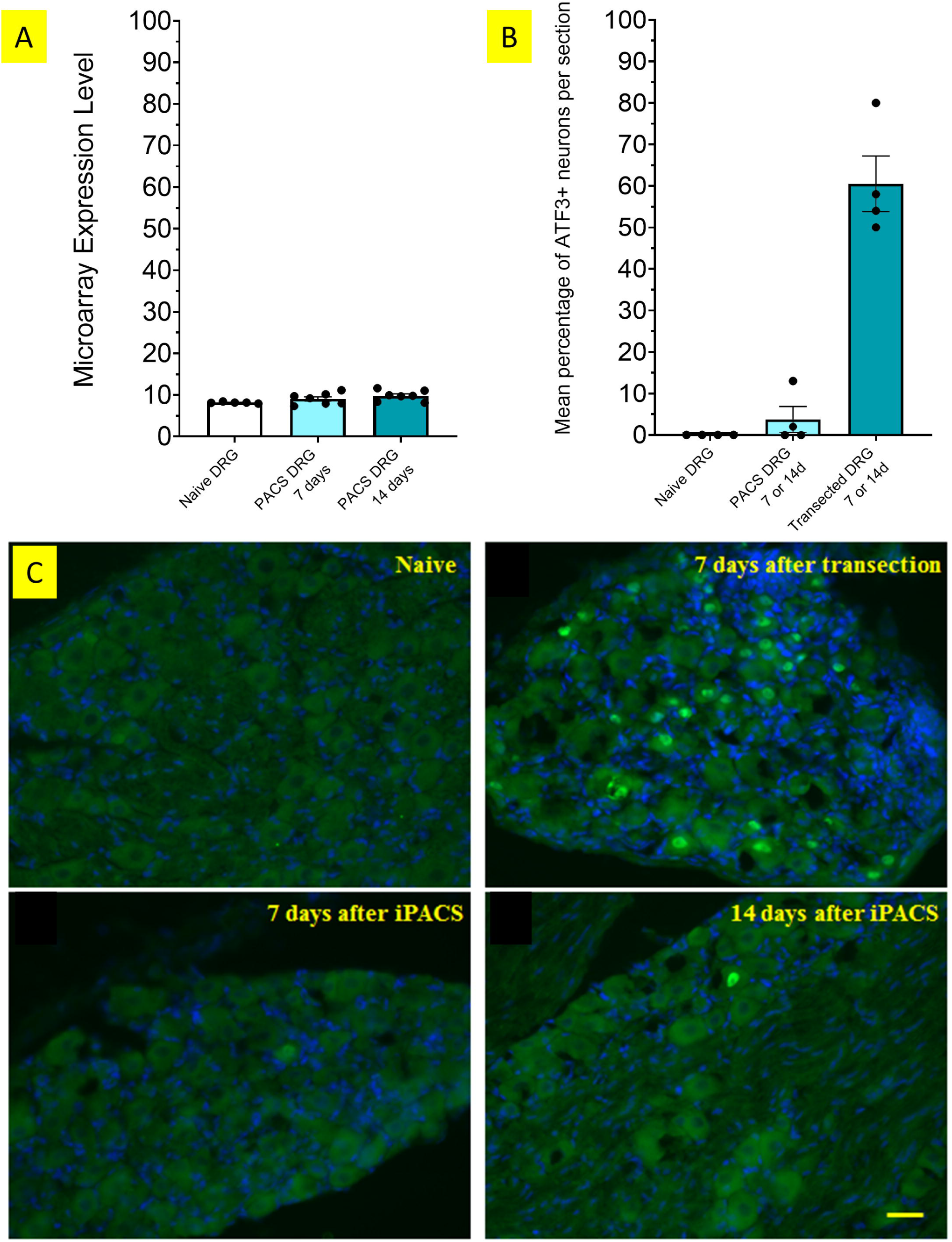
The neuronal injury marker *Activating Transcription Factor 3* (Atf3) was not upregulated by DRG neurons undergoing PACS. (A) Microarray analysis showed that *ATF3* mRNA was expressed at a very low level in naïve T11 DRG (n=5) and that there was no upregulation in T11 DRG undergoing PACS at 7 (n=7) or 14 days after surgery (n=7) (ANOVA F_2,16_=2.8; p>0.05). (B) Very few neurons showed immunoreactivity for ATF3 in naïve T11 DRG or in PACS T11 DRG whereas in adjacent DRG (after injury to their lateral and cutaneous nerves), a large number of neurons showed immunoreactivity for ATF3. Number of DRGs=4/group. (C) Very few neurons showed immunoreactivity for ATF3 in naïve DRG. ATF3 protein was expressed in large numbers of neurons from a positive control tissue (a lumbar DRG whose peripheral branch was transected seven days previously). In contrast, very few neurons showed immunoreactivity for ATF3 in DRG taken either 7 or 14 days after induction of PACS. Images show representative fields of view from each group. Nuclei were counterstained using bisbenzamide (blue). Scale bar, 50 μm.

### Induction of PACS primes neurons for enhanced neurite outgrowth

For cell culture work, successful PACS *in vivo* was confirmed in all rats without exception by “pinch mapping” immediately following spared dermatome surgery (using an indelible pen to mark the denervated zone on the skin; Figure 2C) and then immediately before DRG were harvested for *in vitro* testing (*i*.*e*., at 7 and 14 days; Figure 2D).

To determine a suitable time-point to assess neurite outgrowth, DRG neurons from naive rats were isolated and cultured in defined media for 18 hours (Figure 4A, B) or 24 hours (Figure 4C, D). In line with previous studies of dissociated mature primary DRG neurons (Smith and Skene, 1997; Cafferty et al., 2004), the majority of naive neurons failed to extend many neurites after 18 hr *in vitro* and those that did had short processes (Figure 4B). In comparison, after 24 hr *in vitro* many naive neurons had sprouted with some extending long neurites (Figure 4D).

**Figure 4:**
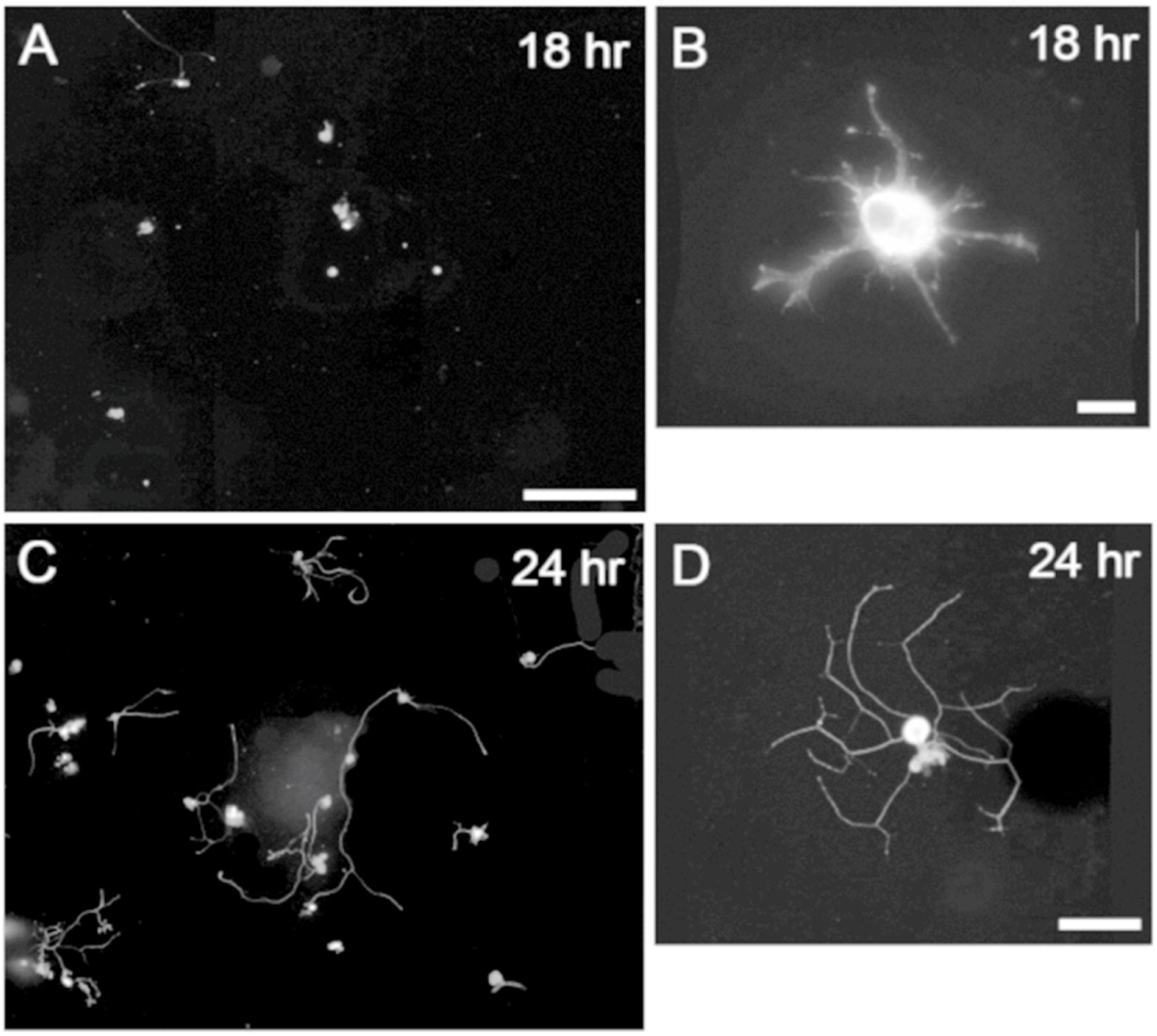
DRG neurons from naïve rats extended neurites poorly in the first 18 hours *in vitro* but arborized by 24 hours. Photomicrographs of dissociated adult DRG neurons from naive animals (A, B) 18 hr and (C, D) 24 hr after initial plating *in vitro*. Cultures were fixed and immunolabelled for βIII tubulin to visualise neurites. Observations revealed less neurite outgrowth after 18 hr where neurons exhibited (A, B) short arborizing neurites compared to the (C, D) long arborizing neurites found after 24 hr of plating. Scale bars: (A, C) 300 µm, (B) 20 µm, (D) 100 µm.

Seven or fourteen days after induction of PACS, neurons from the spared T11 DRG cultured for 18 hours showed increased elongation (Figure 5A) and/or arborization (Figure 5B) relative to naïve neurons (Figure 4). Considering these axon characteristics together, the neurons had longer average neurite lengths than naive neurons (7 days after surgery vs. naïve; *p=*0.03) and 14 days after surgery vs. naïve (*p=*0.04). As expected from prior studies, neurons from DRG with transected nerves also had longer average neurite lengths than naive neurons (7 days after surgery vs. naïve (*p=*0.02) and 14 days after surgery vs. naïve; *p=*0.02; Figure 5C).

**Figure 5:**
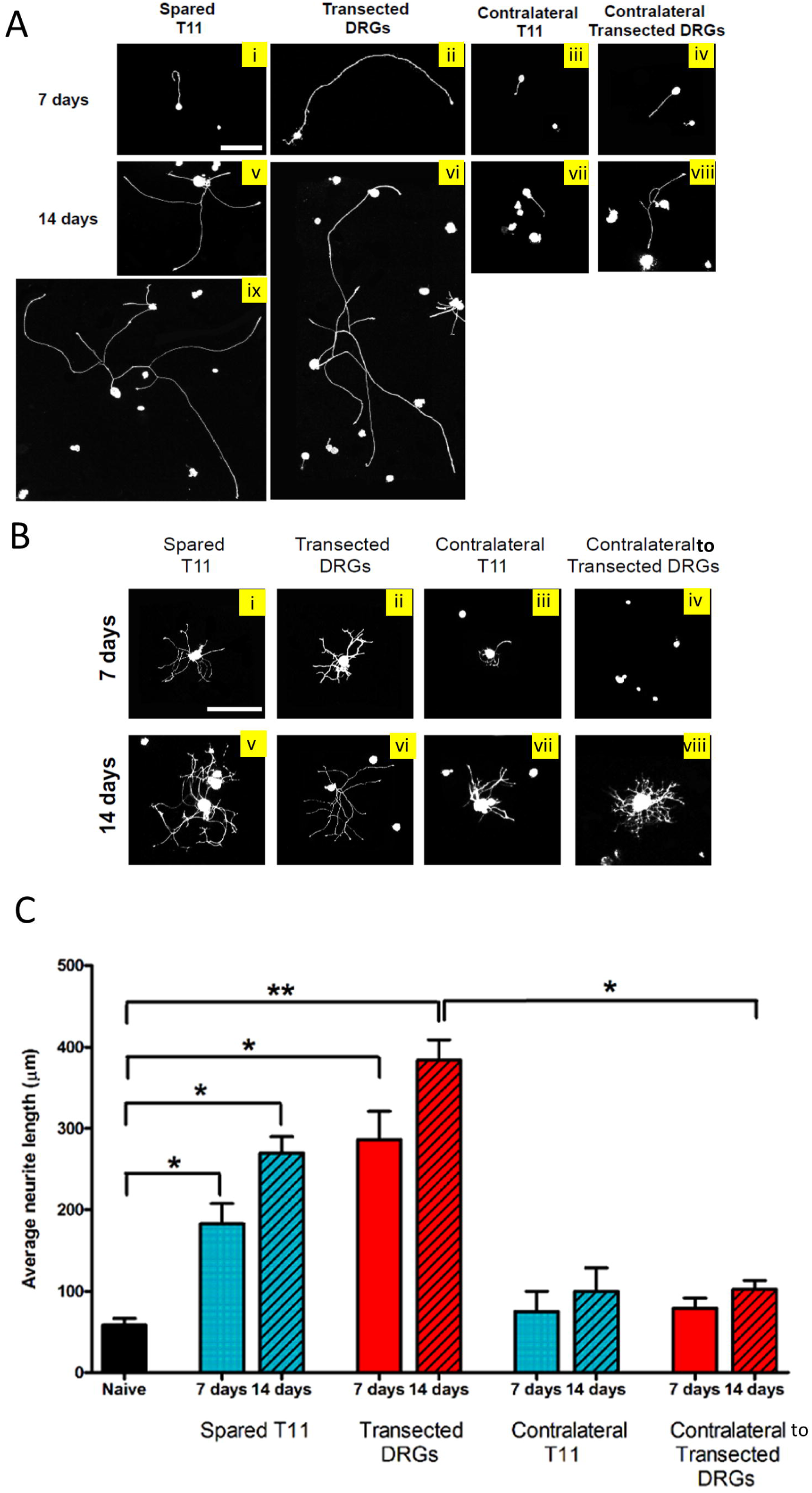
Induction of PACS causes uninjured neurons from T11 DRGs to have longer average neurite lengths than naïve neurons *in vitro*. (A) Photomicrographs show examples of adult DRG neurons from the experimental (i, v, ix), positive control (ii, vi) and negative control groups (iii, iv, vii, viii) elongating after 18 hr in culture. Scale bar, 200 µm. (B) Examples of neurons with neurites classified as “arborizing”. (C) Quantification of average neurite length by Sholl analysis. Results are presented as mean ± SEM and were analysed using Mann Whitney (groups compared to naive neurons) and Wilcoxon (groups compared to contralateral control neurons). Significance is denoted as: * *p* < 0.05, ** *p* < 0.01, *** *p* < 0.001. Naïve *n*=3; 7 days, *n* = 4; 14 days, *n* = 3. After submission, it was found that some images in this Figure were modified digitally to remove cell culture debris in a way that is not appropriate. We intend to replace these with appropriate images once the original files can be obtained.

There was a larger percentage of neurons with a neurite longer than 50 µm in neurons from PACS DRG compared to naïve neurons (naïve vs. 7 days, *p=*0.02; naïve vs. 14 days, *p=*0.03) and from transected DRG compared to naive neurons (naive vs. 7 days, *p=*0.007, naive vs. 14 days, *p=*0.01; Figure 6A).

**Figure 6:**
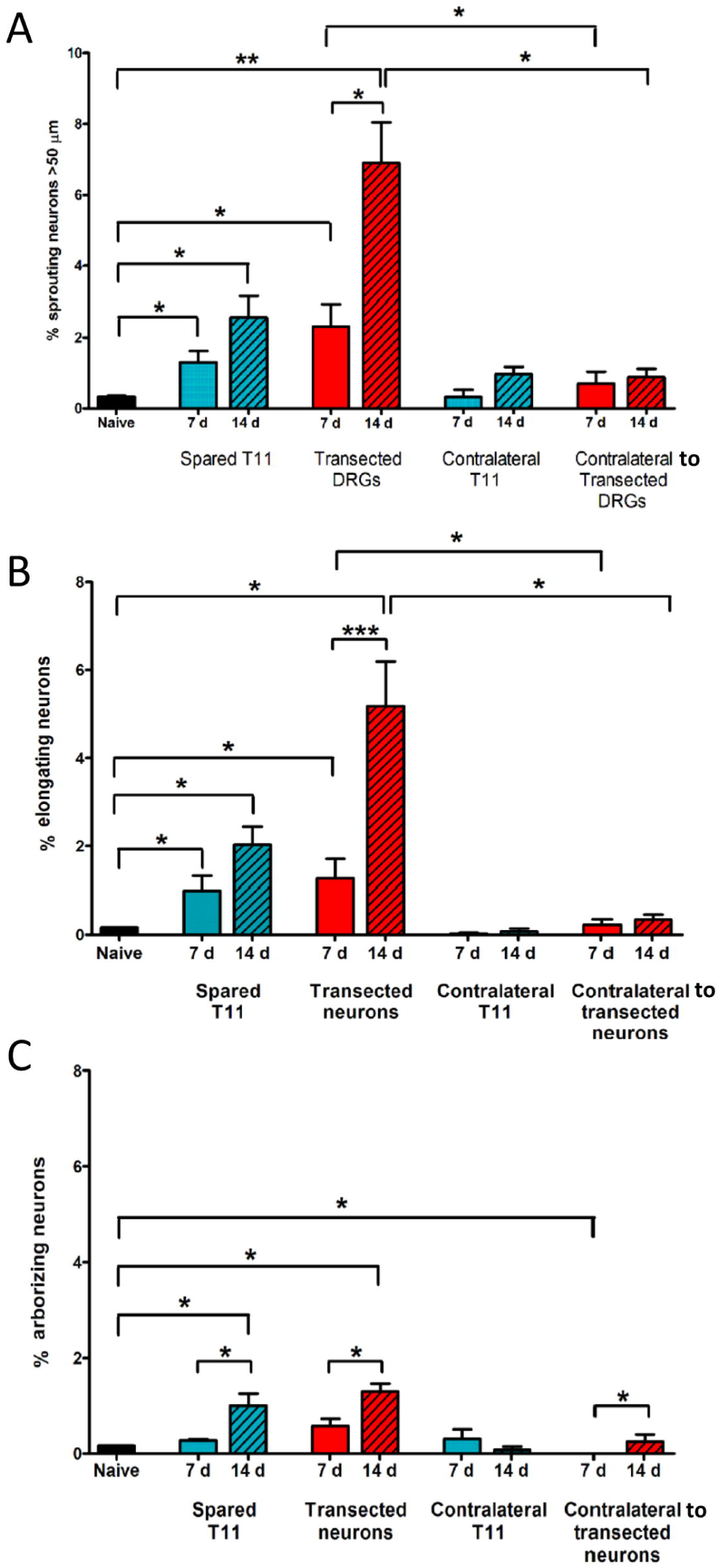
Induction of PACS causes uninjured neurons from T11 DRGs to elongate and/or arborize more than naive contralateral T11 DRG neurons. (A) A larger proportion of neurons from both PACS DRGs and Transected DRGs extended neurites (of either “elongating” or “arborizing” morphology) relative to those from naïve DRGs both 7 and 14 days after induction of PACS. (B) A larger percentage of neurons from both PACS DRGs and Transected DRGs elongated relative to those from naïve DRGs both 7 and 14 days after induction of PACS. (C) A larger percentage of neurons from both PACS DRGs and Transected DRGs arborized relative to those from naïve DRGs only 14 days after induction of PACS. The percentage was larger at 14 days than at 7 days for both PACS DRGs and Transected DRGs. All results are presented as mean ± SEM. Results were analysed using Mann Whitney (groups compared to naive neurons) and Wilcoxon (groups compared to contralateral control neurons). Significance is denoted as: * *p* < 0.05, ** *p* < 0.01, *** *p* < 0.001. Naïve *n*=3; 7 days, *n* = 4; 14 days, *n* = 3.

### A greater proportion percentage of neurons showed arborization after induction of PACS or after transection than in naive DRG

The process of PACS *in vivo* might involve both axonal elongation and arborization to re-innervate large volumes of skin. Using Sholl analysis (Figure 6A), neurons with a mean of <2 branches per 100 µm neurite were categorised as elongating, whereas those with mean >2 branches per 100 µm neurite were categorised as arborizing. Examples of DRG neurons categorised as “elongating” or “arborizing” can be seen in Figures 5A and B, respectively. These two groups were analysed next separately.

A larger percentage of neurons from PACS DRG were categorised as elongating than those from naïve DRG at 7 days (*p*=0.03; Figure 6B) and at 14 days (p=0.04). A larger percentage of neurons from transected DRG were categorised as elongating relative to those from naïve DRG at 7 (*p*=0.05) and 14 days (p=0.02) The percentage of neurons categorised as elongating increased from 7 to 14 days after surgery in transected neurons (*p=*0.001) and showed a strong trend in PACS neurons (*p=*0.07).

A larger percentage of neurons from PACS DRG were categorized as arborizing than those from naive DRG at 14 days (p=0.04; Figure 6C) but not at 7 days (p=0.29). A larger percentage of neurons from transected DRG were categorized as arborizing than those from naive DRG at 14 days (p=0.04) and there was a trend at 7 days (p=0.09).

### Small diameter neurons account for the majority of neurons extending neurites in vitro

We categorised each neuron as having a small diameter (≤35 µm) or a large diameter (>35 µm) *in vitro*. Of the neurons that extended neurites, a greater proportion had a small diameter soma than had a large diameter soma at both seven and fourteen days after induction of PACS (7 days, *p=*0.01; 14 days, *p=*0.05; Figure 7B). There was no difference in the proportion of sprouting neurons with a small diameter between these time points (7 vs. 14 days, *p=*0.2). This same pattern was observed when considering the different characteristics of neurite growth. There was a greater proportion of neurons with elongating axons possessing a small diameter soma than with a large diameter (7 days, *p=*0.02; 14 days, *p=*0.05; Figure 7D), and no difference in proportion between time points (7 vs. 14 days, *p=*0.4). There was a greater proportion of neurons with arborising axons possessing a small diameter soma than with a large diameter (7 days, *p=*0.01; 14 days, *p=*0.05; Figure 7F), and no difference in proportion between time points (7 vs. 14 days, *p=*0.2).

**Figure 7:**
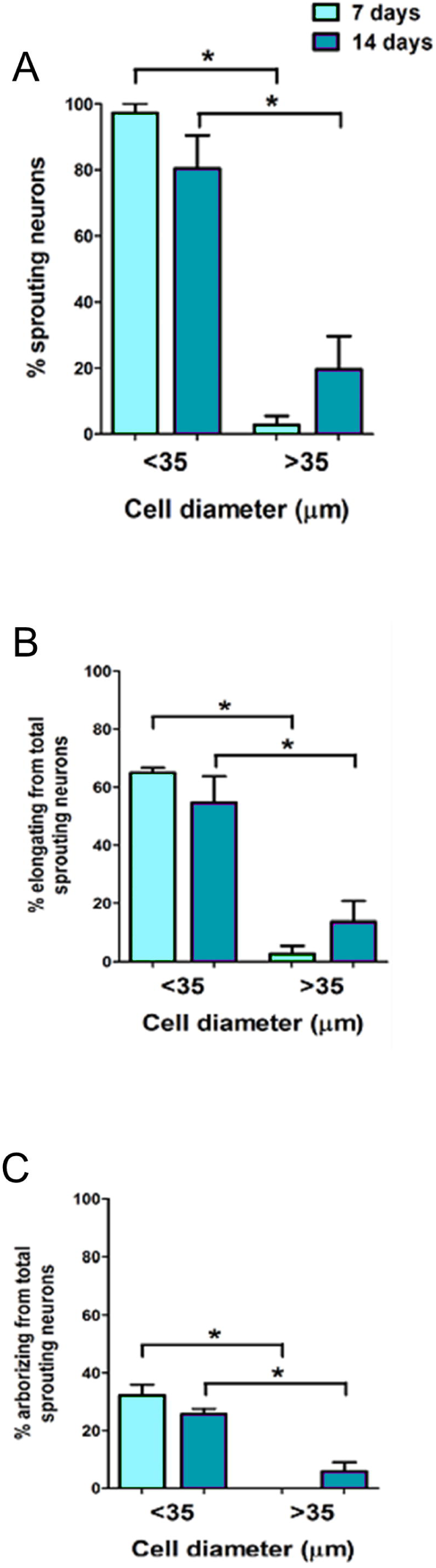
The majority of sprouting neurons had a small diameter cell body. T11 DRG neurons from PACS rats were categorised as small diameter (<35 µm) or larger diameter (>35 µm) sprouting neurons. **(A)** The majority of sprouting neurons were found to be small diameter neurons, including **(B)** elongating and **(C)** arborizing types. Results are presented as mean ± SEM and were analysed using the Wilcoxon test. Significance is denoted as: * *p* < 0.05.

Collectively, these data suggest that the population responsible for the axonal growth observed *in vitro* is the small diameter neurons.

### The majority of sprouting neurons are CGRP negative

Previous studies have reported that Aδ (Nixon et al., 1984) and C fibers from uninjured DRG neurons undergo Primary Afferent Collateral Sprouting *in vivo* (Doucette and Diamond, 1987). Others have shown that conventional conditioning lesions induce a growth state in some but not all types of DRG neurons (Kalous and Keast, 2010). We therefore sought to identify which subtypes of T11 DRG neurons undergo enhanced neurite outgrowth *in vitro* following induction of PACS *in vivo*. We identified sprouting neurons by immunolabeling for beta III tubulin and we co-immunolabelled for the peptide CGRP (Figure 8).

**Figure 8:**
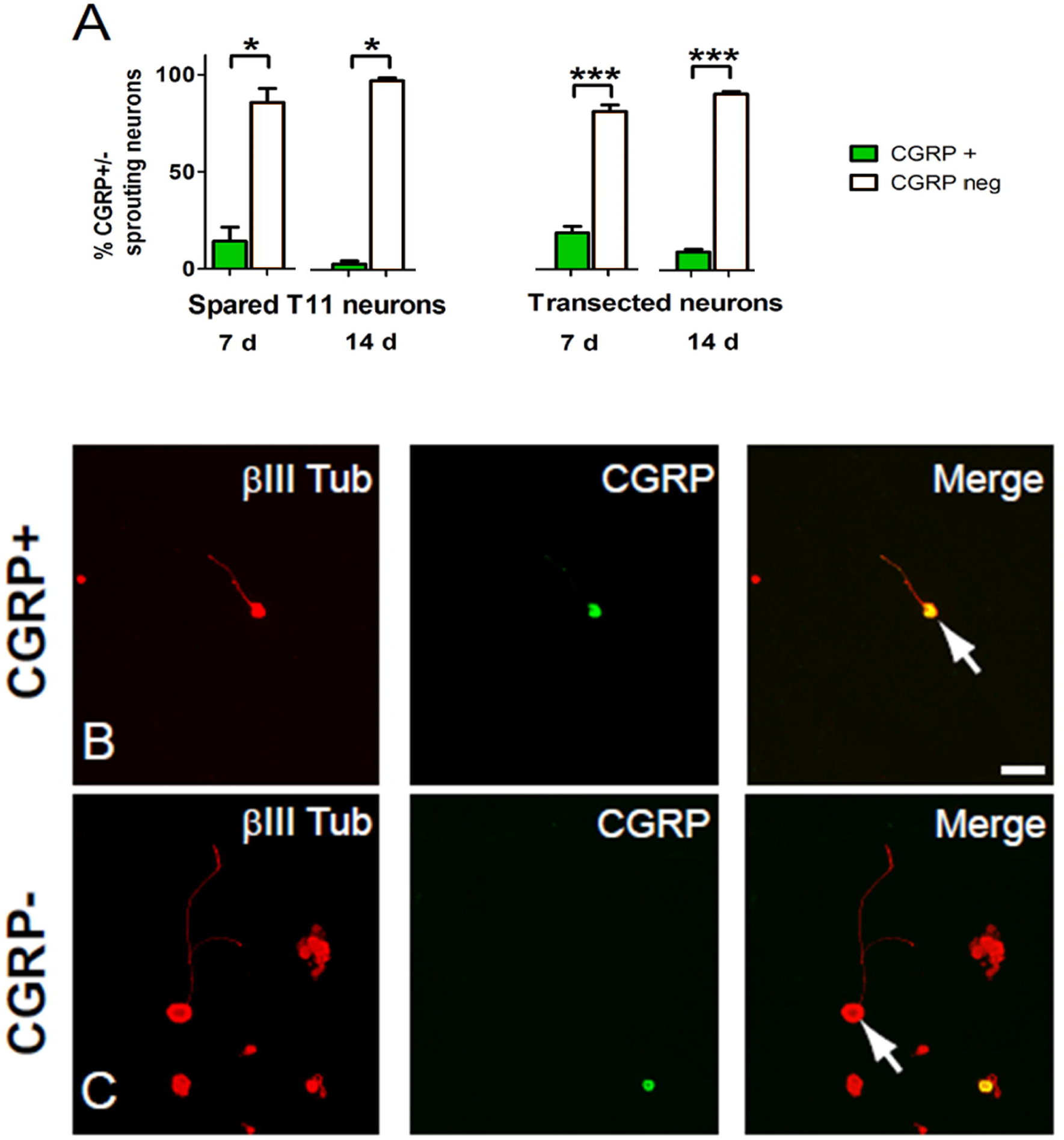
Sprouting neurons are mostly CGRP negative. **(A)** More sprouting neurons from PACS and transected DRGs were CGRP negative than positive. Results are presented as mean ± SEM and were analysed using the Wilcoxon test. Significance is denoted as: * *p* < 0.05, *** *p* < 0.001. **(B-C)** Photomicrographs of dissociated adult β-III-tubulin positive (red) neurons from the uninjured T11 DRG were immunolabelled for CGRP (green) at 14 days after induction of PACS. Scale bar: 50 µm. After submission, it was found that some images in this Figure were modified digitally to remove cell culture debris in a way that is not appropriate. We intend to replace these with appropriate images once the original files can be obtained.

Although a small percentage of T11 DRG neurons extending neurites were CGRP positive, a greater percentage were CGRP negative at both time-points after induction of PACS (Figure 8A-C) (7 days, *p=*0.02; 14 days *p=*0.05). A greater percentage of neurons from DRG with transected nerve were CGRP negative than CGRP positive at 7 days (*p=*0.001) and at 14 days (*p=*0.004) (Figure 7A).

### T11 DRG with neurons undergoing PACS express mRNA for receptors whose ligands are regulated in denervated skin

We have previously performed microarray analysis using RNA from T11 DRG and from denervated skin taken from naïve rats or from rats 7 days or 14 days after induction of PACS *in vivo* (Flight et al., 2014; Harrison et al., 2015; Harrison et al., 2016). Tables 2 and 3 show the genes in DRG and skin (respectively) that were evaluated by real time qRTPCR; results obtained using dChip for microarray analysis are also presented for comparison. As expected, microarray analysis showed that markers of neuronal injury were not upregulated in T11 DRG at 7 or 14 days (*e*.*g*., activating transcription factor 3, Figure 3A; colony stimulating factor 1; p-values>0.05).

**Table 2:** Microarray and real time RT PCR analyses revealed that T11 DRG showed changes in gene expression at 7 and 14 days after induction of PACS. Microarray fold changes are based on dChip estimates of expression values. Some of these results have been presented previously (Harrison et al., 2015). n.s. indicates fold change was not significant. Shading of a box indicates significant change (p<0.05) or a trend.

|  | Microarray |  |  |  |  |  | Real-time RT-PCR |  |  |  |  |
| --- | --- | --- | --- | --- | --- | --- | --- | --- | --- | --- | --- |
| Gene name | Probe set ID | Fold-change |  | P-values |  |  | Fold-change |  | P-values |  |  |
|  |  | 7 days | 14 days | ANOVA | Naive vs.7d | Naive vs.14d | 7 days | 14 days | ANOVA | Naive vs.7d | Naive vs.14d |
| Arl6ip5 | 1369319_at | 1.30 | 1.26 | F(2,16)=4.2;<br>p=0.0347 | t(16)=2.7;<br>p=0.0147 | t(16)=2.4;<br>p=0.0316 | 1.13 | 1.05 | 0.704 | 0.482 | 0.906 |
| BDNF | 1368677_at | 1.66 | 1.77 | F(2,16)=13.3;<br>p=0.0004 | t(16)=4.2;<br>p=0.0007 | t(16)=4.9;<br>p=0.0002 | 1.94 | 1.99 | 0.010 | 0.020 | 0.014 |
|  | 1368678_at | 1.33 | 1.31 | F(2,16)=6.2;<br>p=0.0103 | t(16)=3.2;<br>p=0.0052 | t(16)=3.0;<br>p=0.0084 |  |  |  |  |  |
| Cacna2d1 | 1369649_at | 1.38 | 1.45 | F(2,16)=4.9;<br>p=0.0215 | t(16)=2.5;<br>p=0.022 | t(16)=3.0;<br>p=0.0088 | 2.28 | 3.11 | 0.036 | 0.225 | 0.029 |
| Car8 | 1398431_at | 0.75 | 0.82 | F(2,16)=4.5;<br>p=0.029 | t(16)=3.0;<br>p=0.0094 | t(16)=2.1;<br>p=0.0541 | 0.85 | 0.87 | 0.520 | 0.556 | 0.594 |
| Cdk10 | 1384589_at | 1.28 | 1.22 | F(2,16)=6.6;<br>p=0.0079 | t(16)=3.6;<br>p=0.0026 | t(16)=2.7;<br>p=0.0147 | Not done |  |  |  |  |
|  | 1395122_s_at | 1.3 | 0.96 | F(2,16)=11.7;<br>p=0.0007 | t(16)=3.6;<br>p=0.0022 | t(16)=0.48;<br>p=0.6378 | 0.92 | 0.83 | 0.189 | 0.318 | 0.194 |
| Enpp3 | 1367905_at | 0.65 | 0.83 | F(2,16)=9.4;<br>p=0.002 | t(16)=4.3;<br>p=0.0005 | t(16)=2.1;<br>p=0.0503 | 0.63 | 0.82 | 0.032 | 0.025 | 0.349 |
| Frzb | 1373615_at | 0.94 | 0.71 | F(2,16)=8.5;<br>p=0.003 | t(16)=0.77;<br>p=0.4542 | t(16)=3.7;<br>p=0.0018 | 0.92 | 0.53 | 0.034 | 0.896 | 0.043 |
| Gadd45a | 1368947_at | 1.66 | 1.86 | F(2,16)=4.1;<br>p=0.0353 | t(16)=2.2;<br>p=0.0464 | t(16)=2.8;<br>p=0.0126 | 1.83 | 1.64 | 0.029 | 0.081 | 0.169 |
| GFRα1 | 1387007_at | 1.13 | 1.16 | F(2,16)=3.4;<br>p=0.0579 | t(16)=2.0;<br>p=0.0648 | t(16)=2.5;<br>p=0.0217 | n.s. | n.s. | F(2,17)=0.32;<br>p=0.7287 | t(17)=0.54;<br>p=0.5988 | t(17)=0.22;<br>p=0.8323 |
| GFRα2 | 1369167_at | 0.95 | 0.87 | F(2,16)=3.8;<br>p=0.0453 | t(16)=0.98;<br>p=0.3456 | t(16)=2.7;<br>p=0.0173 | n.s. | n.s. | F(2,16)=1.5;<br>p=0.2526 | t(16)=1.6;<br>p=0.1404 | t(16)=1.5;<br>p=0.145 |
| GFRα3 | 1377680_at | n.s. | n.s. | n.s. | n.s. | n.s. | 0.79 | n.s. | F(2,17)=2.5;<br>p=0.1143 | t(17)=2.2;<br>p=0.0432 | t(17)=1.5;<br>p=0.1462 |
| GFRα4 | 1369730_a_at | n.s. | n.s. | n.s. | n.s. | n.s. | 0.67 | 0.59 | F(2,17)=6.0;<br>p=0.0105 | t(17)=2.7;<br>p=0.0162 | t(17)=3.3;<br>p=0.0042 |
| P75 (Ngfr) | 1368148_at | n.s. | n.s. | n.s. | n.s. | n.s. | n.s. | n.s. | 0.772 | 0.827 | 0.782 |
| Ret b | Various probes | n.s. | n.s. | n.s. | n.s. | n.s. | n.s. | n.s. | F(2,17)=0.57;<br>p=0.5763 | t(17)=0.19;<br>p=0.85 | t(17)=0.78;<br>p=0.4455 |
| Spp1 | 1367581_a_at | 1.46 | 1.70 | F(2,16)=4.1;<br>p=0.0353 | t(16)=2.2;<br>p=0.0464 | t(16)=2.8;<br>p=0.0126 | 1.63 | 1.60 | 0.042 | 0.024 | 0.030 |
| Sema 6a | 1382868_at | 1.57 | 1.48 | F(2,16)=4.9;<br>p=0.0221 | t(16)=3.0;<br>p=0.0088 | t(16)=2.5;<br>p=0.0238 | 1.61 | 1.51 | 0.129 | 0.136 | 0.234 |
| TrkA (Ntrk1) | 1375615_at | n.s. | n.s. | n.s. | n.s. | n.s. | 0.71 | 0.74 | F(2,17)=3.9;<br>p=0.0409 | t(17)=2.6;<br>p=0.0205 | t(17)=2.3;<br>p=0.0336 |
| TrkC (Ntrk3) | 1377308_a_at | 0.82 | 0.80 | F(2,16)=5.2;<br>p=0.0178 | t(16)=2.7;<br>p=0.0156 | t(16)=3.0;<br>p=0.0081 | n.s. | n.s. | F(2,16)=0.1<br>8; p=0.8354 | t(16)=0.57;<br>p=0.5771 | t(16)=0.17;<br>p=0.867 |
|  | 1368939_a_at | n.s. | n.s. | n.s. | n.s. | n.s. | Not done |  |  |  |  |

**Table 3:** Microarray and real time RT PCR analyses revealed that denervated skin showed changes in gene expression at 7 and 14 days after induction of PACS. Microarray fold changes are based on dChip estimates of expression values. Some of these results have been presented previously (Harrison et al., 2015). n.s. indicates fold change was not significant. Shading of a box indicates a significant change (p<0.05). Number of rats per group whose samples passed microarray quality control measures: naïve n=6; day seven n=5; day fourteen n=5.

|  | Microarray |  |  |  |  |  | Real-time RT-PCR |  |  |  |  |
| --- | --- | --- | --- | --- | --- | --- | --- | --- | --- | --- | --- |
| Gene name | Probe set ID | Fold-change |  | P-values |  |  | Fold-change |  | P-values |  |  |
|  |  | 7 days | 14 days | ANOVA | Naive vs.7d | Naive vs.14d | 7 days | 14 days | ANOVA | Naive vs.7d | Naive vs.14d |
| **Artemin** | Not represented on the chip |  |  |  |  |  | 2.32 | 1.62 | <0.001 | <0.001 | 0.048 |
| **GDNF** | 1369598_at | n.s. | n.s. | n.s. | n.s. | n.s. | n.s. | n.s. | F(2,14)=1.3;<br>p=0.3010 | t(14)=1.6;<br>p=0.1369 | t(14)=0.33;<br>p=0.7454 |
| **Neurturin** | 1374048_at | n.s. | n.s. | n.s. | n.s. | n.s. | n.s. | n.s. | F(2,14)=0.4;<br>p=0.6806 | t(14)=0.4;<br>p=0.6952 | t(14)=0.89;<br>p=0.3891 |
| **Ngfb primer pair #01** | 1371259_at | 0.88 | 0.97 | n.s. | t(13)=2.3;<br>p=0.0359 | t(13)=0.52;<br>p=0.6117 | 3.0 | 3.1 | 0.002 | <0.05 | <0.05 |
| **Ngfb primer pair #05** | As above |  |  |  |  |  | 2.6 | 3.0 | 0.001 | <0.05 | <0.05 |
| **NT-3** | 1387267_at | 0.82 | 0.83 | F(2,13)=8.3;<br>p=0.0048 | t(13)=3.6;<br>p=0.031 | t(13)=3.3;<br>p=0.0057 | n.s. | n.s. | 0.499 | 0.717 | 0.493 |
| **Persephin** | 1387533_at | 0.95 | 1.1 | F(2,13)=4.8;<br>p=0.0277 | t(13)=0.95;<br>p=0.3575 | t(13)=2.2;<br>p=0.0476 | 1.65 | 2.14 | F(2,15)=3.9;<br>p=0.042 | t(15)=1.6;<br>p=0.1404 | t(15)=2.8;<br>p=0.0147 |
| **P75 (Ngfr)** | 1368148_at | n.s. | n.s. | n.s. | n.s. | n.s. | 6.54 | n.s. | F(2,15)=6.8;<br>p=0.008 | t(15)=3.7;<br>p=0.0022 | t(15)=1.5;<br>p=0.147 |
| **Ret a** | 1388208_a_at | 0.93 | 1.22 | F(2,13)=11.6;<br>p=0.0013 | t(13)=1.1;<br>p=0.2753 | t(13)=3.7;<br>p=0.0029 | 0.87 | 1.92 | F(2,15)=3.4;<br>p=0.0594 | t(15)=0.31;<br>p=0.7590 | t(15)=2.3;<br>p=0.0377 |
| **Ret b** | Various probes | n.s. | n.s. | n.s. | n.s. | n.s. | n.s. | n.s. | F(2,14)=2.4;<br>p=0.1295 | t(14)=0.51;<br>p=0.6205 | t(14)=1.7;<br>p=0.1082 |

We were interested to determine whether there were changes in expression level in DRG for receptors for ligands whose levels were altered in denervated skin, particularly those known to be expressed by small diameter neurons that sprout in response to induction of PACS *in vivo* and *in vitro* (including NGF and GDNF family receptors and ligands). Others have shown that NGF mRNA is increased in skin, for example (Mearow et al 1993). Surprisingly, our microarray analysis did not detect upregulation of Nerve Growth Factor mRNA in skin (NGF beta probe set 1371259_at; NGF gamma probe set 1367961_at); however, qRTPCR using two different primer pairs (Table 1) confirmed that NGF beta mRNA was upregulated (∼3-fold) in skin at 7 days and 14 days (Figure 9; Table 3). Although microarray analysis did not detect changes in TrkA in DRG (Table 2), qRTPCR identified decreases at day 7 and 14 (Figure 10; Table 3). qRTPCR showed that p75NTR was also upregulated in skin at 7 but not 14 days after induction of PACS (Figure 9; Table 3) but not in DRG at 7 or 14 days (Table 2).

**Figure 9:**
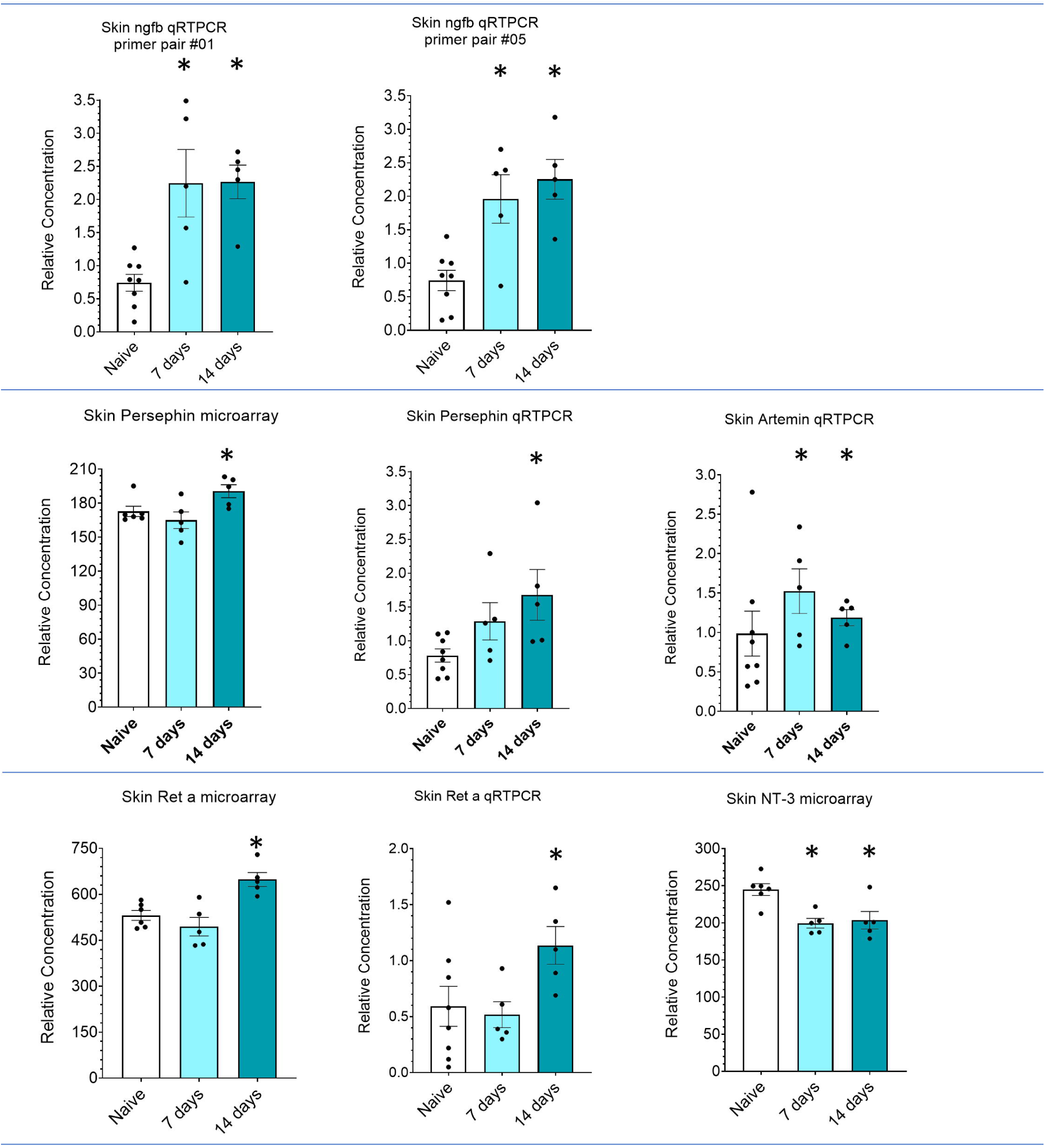
The level of expression of growth factor ligands were regulated in denervated skin. Graphs show microarray or qRTPCR data presented as mean ± SEM. * indicates *p* < 0.05 by Fisher’s LSD test protected by a preceding significant one-way ANOVA (*p* < 0.05). Number of rats for microarray experiment: naïve n=6, 7 days n=5, 14 days n=5. Number of rats for qRTPCR experiment: naïve n=8, 7 days n=5, 14 days n=5. F values and degrees of freedom for these data can be found in Table 2. Primer information is in Table 1.

**Figure 10:**
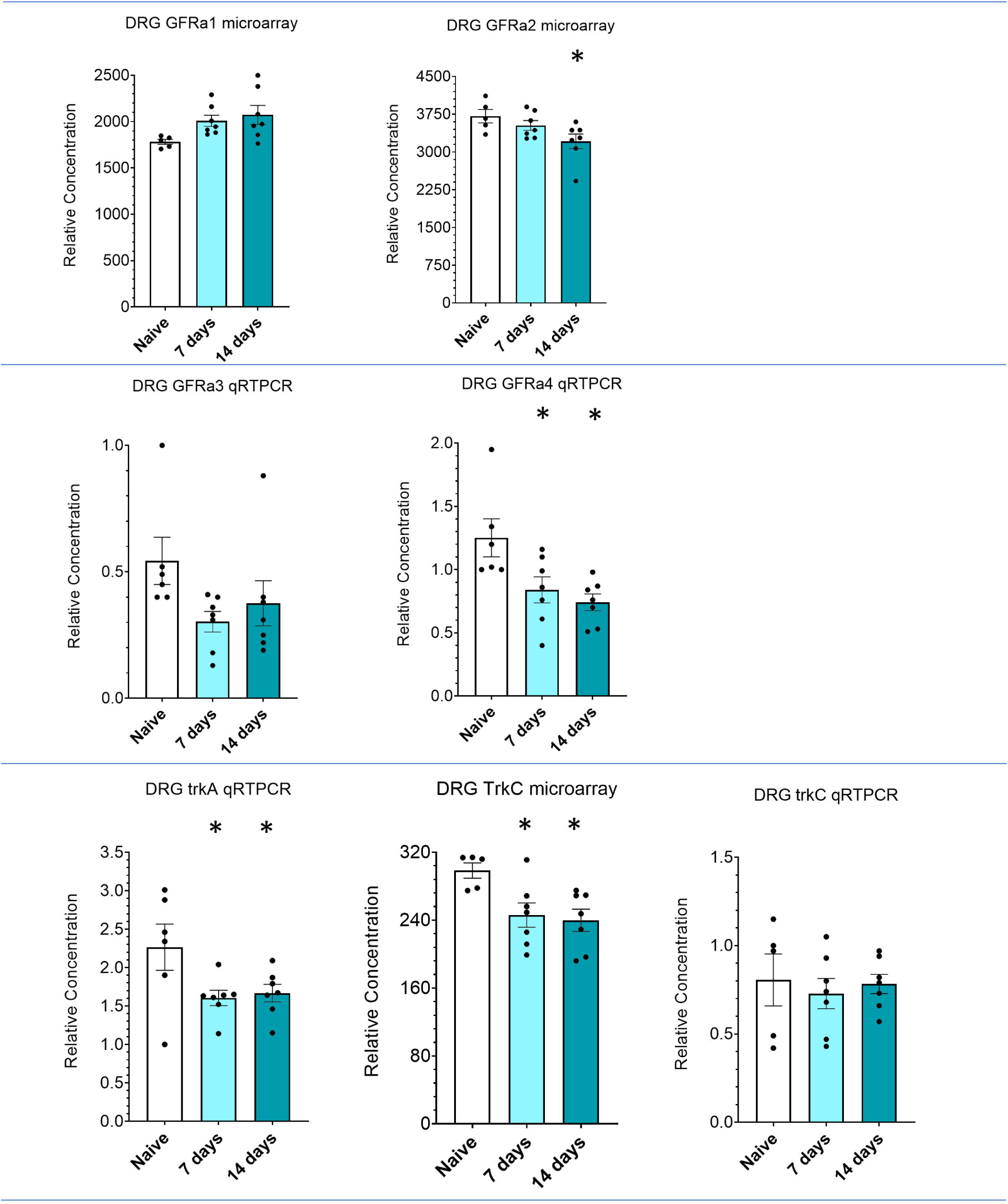
The level of expression of growth factor receptors were regulated in T11 DRG undergoing PACS. Graphs show microarray or qRTPCR data presented as mean ± SEM. * indicates *p* < 0.05 by Fisher’s LSD test protected by a preceding significant one-way ANOVA (*p* < 0.05). Number of rats for microarray experiment: naïve n=5, 7 days n=7, 14 days n=7. Number of rats for qRTPCR experiment: naïve n=6, 7 days n=7, 14 days n=7. F values and degrees of freedom for these data can be found in Table 3. Primer information is in Table 1.

Small diameter DRG neurons are also known to express receptors for GDNF family receptor alpha (GFRα) 1, 2, 3 and 4 and c-ret (Bennett et al., 1998; Bennett et al 2000). Interestingly, the GDNF family ligand, Persephin, was shown by microarray analysis and qRTPCR to be regulated in skin at 14 days after induction of PACS (Figure 9; Table 3). qRTPCR showed that Artemin, another GDNF family ligand, was upregulated in skin at 7d and 14 days after induction of PACS (Figure 9; Table 3) [*n*.*b*., Artemin was not represented by a probe set on the microarray chips]. There was no evidence that GDNF or Neurturin was regulated in skin by either microarray analysis or qRTPCR (Table 3). Regarding receptors, qRTPCR found evidence for downregulation in DRG of GFRα4 at 7 and 14 days (Figure 10; Table 2) although this was not detected by microarray analysis. Microarray analysis indicated slight upregulation of GFRα2 in DRG at 14 days but this was not validated by qRTPCR (Figure 10; Table 2). There was no strong evidence for regulation of GFRα1 or GFRα3 or Ret isoform b by either method in DRG but there was evidence that Ret isoform a was upregulated in skin at 14 days by both microarray and qRTPCR (Figure 9; Table 3).

Large diameter DRG neurons serving low-threshold mechanoreception functions in the skin express the TrkC receptor for neurotrophin-3 (NT-3; McMahon et al 1994; Krimm et al., 2000) and the available evidence indicates that these axons do not sprout during PACS (Jackson and Diamond, 1983, 1984; Horch 1981). Consistent with this, microarray analysis showed evidence for slight downregulation of NT-3 in skin (−1.2 fold) at both 7 days and 14 days (Figure 9; Table 3) although qRTPCR did not confirm any regulation; above all, there was no evidence for upregulation of NT-3 in skin and this may account for the failure of sprouting of TrkC expressing DRG neurons during PACS. Microarray analysis also showed evidence for downregulation of TrkC in DRG by 1.2 fold at both 7 days and 14 days (Table 2) although qRTPCR did not confirm this.

Finally, qRTPCR validated other changes in gene expression in T11 DRG at 7 and 14 days after induction of PACS including BDNF, CCL2, Frzb, Grhl3, Spp1 (osteopontin) and Enpp3, Cacna2d1 and Gadd45a (Table 2). Future work may determine whether these genes are involved in priming of DRG neurons for outgrowth once dissociated and maintained in defined media lacking neurotrophins and growth factors *in vitro*.

## Discussion

Adult mammalian sensory neurons are capable either of axonal regeneration of cut axons or of collateral sprouting of non-injured axons induced by various stimuli including loss of neighbouring innervation (Mahar and Cavalli, 2018; Lorenzana et al., 2015). Prior injury (“conditioning”) is known to induce an enhanced axonal outgrowth when sensory neurons are subsequently grown *in vitro* (Smith & Skene, 1990). It was not known if the changes induced by collateral sprouting *in vivo* may prime uninjured neurons for enhanced growth *in vitro*.

As expected, the transection of multiple thoracic cutaneous nerves either side of the uninjured T11 nerves produced a CTM-responsive region isolated by areas that initially did not exhibit CTM reflex responses. The responsive region expanded over 7 to 14 days into previously unresponsive areas (Jackson and Diamond, 1981; Nixon et al., 1984; Diamond et al., 1987; Doucette and Diamond, 1987; Diamond et al., 1992b; Pertens et al., 1999): this confirms that uninjured axons from the spared T11 DRG sprouted and restored functionally useful connections (Isaacson and Crutcher, 1998; Jackson and Diamond, 1981; Doucette and Diamond, 1987; Jackson and Diamond, 1984). Further, we confirmed that few if any neurons undergoing PACS were injured during surgery by immunolabeling for an injury-related protein (ATF3) (Tsujino et al 2000).

### Induction of PACS primed DRG neurons for axonal elongation or arborization in vitro

DRG neurons were dissociated seven or fourteen days after induction of PACS and grown for 18h in defined media lacking growth factors or neurotrophins. A larger proportion of PACS neurons extended neurites than naive neurons, including some with either elongating or arborizing morphologies. Others had previously shown that a conditioning lesion causes injured neurons to initiate extension of neurites more rapidly in vitro and that neurites tend to elongate rather than arborize (Kalous and Keast, 2010; Cafferty et al., 2004; Cafferty et al., 2001; Smith and Skene, 1997). Our experiments extend these results and show that an accelerated growth process can also be initiated without injury by induction of collateral sprouting *in vivo*.

Although the absolute proportion of PACS neurons that sprouted may appear small, it is much larger when considered relative to the subpopulation that would be affected in this surgical model. This model does not denervate the entire dermatome of the neighboring segments: only the dorsal and lateral cutaneous nerves were transected, leaving the ventral cutaneous nerves intact. Thus, only those neurons with axons innervating that portion of the T11 dermatome served by the dorsal cutaneous nerve and lateral cutaneous nerve (but not ventral cutaneous nerve) would be subjected to a PACS-inducing environment. This is in the range of only 40% of the DRG neurons in the rat T11 DRG (Hill et al., 2010). The remainder of the T11 DRG serves targets not adjacent to denervated tissue in the PACS model, including viscera and muscles (which therefore one would expect not to sprout). Approximately 5% of the total number of T11 PACS DRG neurons displayed axonal outgrowth. This 5% of the total population is 12.5% of the total sub-population exposed to a PACS-inducing environment, as defined by neurons with axons in the dorsal or lateral cutaneous nerves. Of those DRG neurons projecting axons through the dorsal or lateral cutaneous nerves, some subtypes are known to not undergo PACS (see below), and others may project to a central portion of the T11 dermatome which is likely to be less-influenced by denervation of the neighboring dermatomes. In summary, our results show that a significant proportion of PACS-primable uninjured neurons underwent enhanced growth *in vitro*.

### Phenotype of neurons that sprouted in vitro after induction of PACS

Conditioning lesions induce a growth state in some but not all types of injured DRG neurons (Kalous and Keast, 2010). We examined the phenotype of uninjured neurons that are primed by PACS. Previous studies have reported that Aδ (Nixon et al., 1984) and C fibers from uninjured DRG neurons undergo Primary Afferent Collateral Sprouting *in vivo*, in contrast to large-diameter A-fiber neurons (Doucette and Diamond, 1987; Horch 1981). We found that the majority of neurons sprouting after PACS were small-diameter neurons which include unmyelinated C fibers and likely also myelinated Aδ fibers. Immunolabeling revealed that some neurons undergoing sprouting were CGRP-positive but that the majority were CGRP-negative. Considering these each in turn:

#### CGRP-positive neurons

These TrkA-positive afferents are capable of robust PACS that is dependent on NGF (Diamond et al., 1992b; Diamond et al., 1987; Doucette and Diamond, 1987; Mearow et al., 1993; Pertens et al., 1999). It is therefore not unexpected that these neurons should be affected by the in vivo PACS condition.

#### CGRP-negative neurons

It is less predictable that these neurons might be affected by PACS, as they have not previously been reported to undergo PACS, though this has never been ruled out. **1)** One possibility is that CGRP-positive neurons secrete a factor within the DRG that primes its CGRP-negative neighbours: candidates might include BDNF. **3)** Another possibility is that CGRP-negative neurons do not undergo PACS *in vivo* but exposure to the denervated skin primes them for subsequent growth *in vitro*. **3)** The third possibility is that CGRP-negative neurons undergo PACS *in vivo* which then primes them for growth *in vitro*. However, if CGRP-negative neurons do undergo PACS *in vivo* then they have been overlooked to date, which is entirely possible for the following reasons: **a)** Silver staining indicated that collateral sprouting is halted by anti-NGF treatments that could be expected to affect only the TrkA-positive, CGRP-positive neurons (Diamond et al., 1992b), presuming a direct effect only. However, these silver stains would not have detected any CGRP-negative, unmyelinated axons because the method preferentially labels neurofilament proteins which are largely absent in unmyelinated axons but which are present in Aδ axons (many of which are trkA+/CGRP+) (Gambetti et al., 1981; Phillips et al., 1983; Dahl et al., 1981). **b)** Moreover, if CGRP-negative neurons do undergo PACS *in vivo* then they may have been missed by the CTM reflex readout: it is the TrkA-positive (CGRP-positive) neurons which drive the CTM reflex in uninjured rats. Evidence for this is that the CTM reflex is absent in rats where TrkA-positive neurons have been ablated using a neurotoxin which spared other small neurons (binding IB4 or being positive for somatostatin or P2×3) (Petruska et al., 2014).

What, then, is the phenotype of the CGRP-negative neurons that sprout *in vitro*? The CGRP-negative population is often characterized by their binding of the lectin IB4, but this binding is rapidly lost upon dissociation and culturing unless growth factors are added (Averill et al., 2004), precluding our use of IB4-binding (given our choice of media). We propose that these CGRP-negative small-diameter neurons are almost certainly cutaneous (as most visceral and muscular small-diameter neurons express CGRP (e.g., Bennett et al., 1996), and are therefore likely in the sub-population exposed to a PACS-inducing environment by the spared dermatome surgery. It is possible that some of the small-diameter, CGRP-negative neurons that sprout *in vitro* express GDNF receptors (Bennett et al., 1998; Bennett et al., 2000) because, as described below, ligands for these receptors are increased in the denervated skin. Future experiments could address this possibility directly.

### Induction of PACS alters gene expression in DRG and in denervated skin

We previously used microarray analysis to identify genes that are differentially regulated in uninjured T11 DRG housing neurons undergoing PACS (Harrison et al., 2015; Harrison et al., 2016) and in the corresponding denervated skin (Flight et al,. 2014). In comparison to naive tissue, significant differences in gene expression were found in hundreds of genes at 7 and 14 days after surgery in both PACS DRGs and denervated skin. More genes and biological processes achieved statistical significance at 14 days after surgery than 7 days after surgery. Might exposure of uninjured DRG neurons to secreted factors in the denervated skin prime their intrinsic transcriptional program for enhanced axon growth that persists after being explanted and grown in media lacking neurotrophins and growth factors?

Genes that were upregulated in the skin included NGF. This supports prior work (Mearow et al 1993) and it may explain the sprouting of TrkA-positive axons. Other genes that were upregulated in skin included the GDNF family ligands Persephin and Artemin. This is interesting because some small-diameter DRG neurons are known to express receptors for these and other GDNF family ligands including GDNF receptor alpha (GFRα) 1, 2, 3 and 4 and c-ret (Bennett et al., 1999; Bennett et al., 2000). It is possible therefore that these GFRα-positive (TrkA-negative) neurons sprout during PACS (although see comments above). Future work may examine whether neurons expressing GFRα receptors undergo PACS (by responding to GDNF family ligands) and become primed for enhanced outgrowth *in vitro*.

The role of target-derived factors in the priming effect we have described merits further study in the future. It is possible that the PACS environment triggers an intrinsic change in the DRG neurons which is carried into the explanted assay system independent of the presence of skin-derived factors, a possibility supported by the numerous gene expression changes documented in DRG in our microarray study. Alternatively, the enhanced growth state may be due to the expression of a gene program enabling receptivity to extrinsic factors, and the presence of the extrinsic factors is then required. In this case, the priming we observe may be transient, in that the increased rate of outgrowth is not sustained beyond 18h of explantation (i.e., after removal from the skin-derived extrinsic factors and induction of injury).

### Causal control over Primary Afferent Collateral Sprouting

Finally, Primary Afferent Collateral Sprouting can be inhibited or enhanced by targeting some of the molecules identified in our microarray screen. For example, daily administration of antisera against beta NGF halts Primary Afferent Collateral Sprouting into denervated skin (Diamond et al 1987) whereas intradermal injections of NGF induce collateral sprouting *de novo* (Diamond et al 1992b). We previously showed that induction of PACS causes upregulation of CD2-associated protein (CD2AP) in DRG and that, in PC12 cells, overexpression of CD2AP enhances neurite outgrowth whilst siRNA-mediated knockdown of CD2AP reduces neurite outgrowth (Harrison et al 2016). In the future, conditional deletion of NGF or GDNF family ligands in skin (or their receptors in DRG) could be performed after induction of PACS. Alternatively, these ligands might be supplemented to facilitate axon growth and functional recovery after damage to skin.

## Conclusions

The findings presented here provide evidence that induction of PACS *in vivo* activates a growth response in uninjured neurons which enhances neurite outgrowth after subsequent dissociation and culture, constituting a novel form of priming. The enhanced neurite outgrowth includes axonal elongation and/or arborization. We also identified some genes which may be involved in the PACS-induced enhancement of axon elongation or arborization.

## Acknowledgments

Thanks to Dr. Ben Harrison for providing the original raw data for Figure 2E. Yunhee Lee and Rebecca Hall performed real time qRTPCR. This work was funded by a grant from the MRC (G0600998) to Dr. Moon, and from the National Institutes of Health (NS094741) to Dr. Petruska. The funding agencies had no involvement in the planning or execution of the study. All three authors designed the research, JP and SS did surgeries, SS analysed data and all three authors contributed to writing the paper.

The authors declare that no conflict of interest exists.

## Data Accessibility

The microarray data may be found at the Gene Expression Omnibus (GEO) with accession numbers GSE72551 (DRG) and GSE54356 (skin). The data for qRTPCR and histology images are available upon request. The cell culture primary data is no longer available due to loss of the hard drive.

## Reference List

Abbadie C, Basbaum AI (1998) The contribution of capsaicin-sensitive afferents to the dorsal root ganglion sprouting of sympathetic axons after peripheral nerve injury in the rat. Neurosci Lett 253:143–146.

Apfel SC, Wright DE, Wiideman AM, Dormia C, Snider WD, Kessler JA (1996) Nerve growth factor regulates the expression of brain-derived neurotrophic factor mRNA in the peripheral nervous system. Mol Cell Neurosci 7:134–142.

Aszmann OC, Muse V, Dellon AL (1996) Evidence in support of collateral sprouting after sensory nerve resection. Ann Plast Surg 37:520–525.

Averill S, Michael GJ, Shortland PJ, Leavesley RC, King VR, Bradbury EJ, McMahon SB, Priestley JV (2004) NGF and GDNF ameliorate the increase in ATF3 expression which occurs in dorsal root ganglion cells in response to peripheral nerve injury. Eur J Neurosci 19:1437–45.

Bennett DL, Dmietrieva N, Priestley JV, Clary D, McMahon SB (1996) trkA, CGRP and IB4 expression in retrogradely labelled cutaneous and visceral primary sensory neurones in the rat. Neurosci Lett 206:33–36.

Bennett DL, Michael GJ, Ramachandran N, Munson JB, Averill S, Yan Q, McMahon SB, Priestley JV (1998) A distinct subgroup of small DRG cells express GDNF receptor components and GDNF is protective for these neurons after nerve injury. J Neurosci 18:3059–3072.

Bennett DL, Boucher TJ, Armanini MP, Poulsen KT, Michael GJ, Priestley JV, Phillips HS, McMahon SB, Shelton DL. 2000 The Glial Cell Line-Derived Neurotrophic Factor Family Receptor Components Are Differentially Regulated within Sensory Neurons after Nerve Injury. The Journal of Neuroscience, 2000, 20(1):427–437.

Bisby MA, Tetzlaff W, Brown MC (1996) GAP-43 mRNA in mouse motoneurons undergoing axonal sprouting in response to muscle paralysis of partial denervation. Eur J Neurosci 8:1240–1248.

Blits, B., Oudega, M., Boer, G.J., Bartlett Bunge, M., Verhaagen, J., 2003. Adeno-associated viral vector-mediated neurotrophin gene transfer in the injured adult rat spinal cord improves hind-limb function. Neuroscience 118, 271–281.

Braz JM, Basbaum AI (2009) Triggering genetically-expressed transneuronal tracers by peripheral axotomy reveals convergent and segregated sensory neuron-spinal cord connectivity. Neuroscience 163:1220–1232.

Brenan A (1986) Collateral reinnervation of skin by C-fibres following nerve injury in the rat. Brain Res 385:152–155.

Brown MC, Holland RL, Hopkins WG (1981) Motor nerve sprouting. Annu Rev Neurosci 4:17–42.

Cafferty WB, Gardiner NJ, Das P, Qiu J, McMahon SB, Thompson SW (2004) Conditioning injury-induced spinal axon regeneration fails in interleukin-6 knock-out mice. J Neurosci 24:4432–4443.

Cafferty WB, Gardiner NJ, Gavazzi I, Powell J, McMahon SB, Heath JK, Munson J, Cohen J, Thompson SW (2001) Leukemia inhibitory factor determines the growth status of injured adult sensory neurons. J Neurosci 21:7161–7170.

Cho HJ, Kim JK, Park HC, Kim JK, Kim DS, Ha SO, Hong HS (1998) Changes in brain-derived neurotrophic factor immunoreactivity in rat dorsal root ganglia, spinal cord, and gracile nuclei following cut or crush injuries. Exp Neurol 154:224–230.

Cotman CW, Nieto-Sampedro M, Harris EW (1981) Synapse replacement in the nervous system of adult vertebrates. Physiol Rev 61:684–784.

Dahl D, Bignami A, Bich NT, Chi NH (1981) Immunohistochemical localization of the 150K neurofilament protein in the rat and the rabbit. J Comp Neurol 195:659–666.

Davis BM, Goodness TP, Soria A, Albers KM (1998) Over-expression of NGF in skin causes formation of novel sympathetic projections to trkA-positive sensory neurons. Neuroreport 9:1103–1107.

Devor M, Schonfeld D, Seltzer Z, Wall PD (1979) Two modes of cutaneous reinnervation following peripheral nerve injury. J Comp Neurol 185:211–220.

Diamond J, Cooper E, Turner C, Macintyre L (1976) Trophic regulation of nerve sprouting. Science 193:371–377.

Diamond J, Coughlin M, Macintyre L, Holmes M, Visheau B (1987) Evidence that endogenous beta nerve growth factor is responsible for the collateral sprouting, but not the regeneration, of nociceptive axons in adult rats. Proc Natl Acad Sci U S A 84:6596–6600.

Diamond J, Foerster A, Holmes M, Coughlin M (1992a) Sensory nerves in adult rats regenerate and restore sensory function to the skin independently of endogenous NGF. J Neurosci 12:1467–1476.

Diamond J, Holmes M, Coughlin M (1992b) Endogenous NGF and nerve impulses regulate the collateral sprouting of sensory axons in the skin of the adult rat. J Neurosci 12:1454–1466.

Doubleday B, Robinson PP (1994) Nerve growth factor depletion reduces collateral sprouting of cutaneous mechanoreceptive and tooth-pulp axons in ferrets. J Physiol 481 (Pt 3):709–718.

Doucette R, Diamond J (1987) Normal and precocious sprouting of heat nociceptors in the skin of adult rats. J Comp Neurol 261:592–603.

Ernfors P, Rosario CM, Merlio JP, Grant G, Aldskogius H, Persson H (1993) Expression of mRNAs for neurotrophin receptors in the dorsal root ganglion and spinal cord during development and following peripheral or central axotomy. Brain Res Mol Brain Res 17:217–226.

Fitzgerald M (1985) The sprouting of saphenous nerve terminals in the spinal cord following early postnatal sciatic nerve section in the rat. J Comp Neurol 240:407–413.

Flight RM, Harrison BJ, Mohammad F, Bunge MB, Moon, LDF, Petruska JC, Rouchka EC (2014) categoryCompare, an analytical tool based on feature annotations. Front. Genet. 5:98.

Gallo G, Lefcort FB, Letourneau PC (1997) The trkA receptor mediates growth cone turning toward a localized source of nerve growth factor. J Neurosci 17:5445–5454.

Gambetti P, Autilio GL, Papasozomenos SC (1981) Bodians silver method stains neurofilament polypeptides. Science 213:1521–1522.

Gavazzi I, Kumar RD, McMahon SB, Cohen J (1999) Growth responses of different subpopulations of adult sensory neurons to neurotrophic factors in vitro. Eur J Neurosci 11:3405–3414.

Genc B, Ulupinar E, Erzurumlu RS (2005) Differential Trk expression in explant and dissociated trigeminal ganglion cell cultures. J Neurobiol 64:145–156.

Giehl KM, Tetzlaff W (1996) BDNF and NT-3, but not NGF, prevent axotomy-induced death of rat corticospinal neurons in vivo. Eur J Neurosci 8:1167–1175.

Gloster A, Diamond J (1992) Sympathetic nerves in adult rats regenerate normally and restore pilomotor function during an anti-NGF treatment that prevents their collateral sprouting. J Comp Neurol 326:363–374.

Gloster A, Diamond J (1995) NGF-dependent and NGF-independent recovery of sympathetic function after chemical sympathectomy with 6-hydroxydopamine. J Comp Neurol 359:586–594.

Goodman LJ, Valverde J, Lim F, Geschwind MD, Federoff HJ, Geller AI, Hefti F (1996) Regulated release and polarized localization of brain-derived neurotrophic factor in hippocampal neurons. Mol Cell Neurosci 7:222–238.

Gutmann, E, 1942. Factors Affecting Recovery of Motor Function after Nerve Lesions. J Neurol Psychiatry 5, 81–95.

Harrison BJ, Venkat G, Hutson TH, Rau KK, Bunge MB, Mendell LM, Gage FH, Johnson RD, Hill CE, Rouchka EC, Moon LDF., Petruska JC, 2015. Transcriptional changes in sensory ganglion associated with primary afferent axon collateral sprouting in spared dermatome model. Genomics Data 6, 249–252.

Harrison BJ, Venkat G, Lamb J, Hutson TH, Drury C, Hill CE, Rouchka EC, Moon LDF, Petruska JC, 2016. The adapter protein CD2AP is a coordinator of neurotrophin signalling-mediated axon arbour plasticity. J Neurosci 36, 4259–4275.

Haubensak W, Narz F, Heumann R, Lessmann V (1998) BDNF-GFP containing secretory granules are localized in the vicinity of synaptic junctions of cultured cortical neurons. J Cell Sci 111 (Pt 11):1483–1493.

Healy C, LeQuesne PM, Lynn B (1996) Collateral sprouting of cutaneous nerves in man. Brain 119 (Pt 6):2063–2072.

Hill, CE, Harrison, BJ, Rau, KK, Hougland, MT, Bunge, MB, Mendell, LM, Petruska, JC, 2010. Skin incision induces expression of axonal regeneration-related genes in adult rat spinal sensory neurons. J Pain 11, 1066–1073.

Horch K (1981) Absence of functional collateral sprouting of mechanoreceptor axons into denervated areas of mammalian skin. Exp Neurol 74:313–317.

Hu-Tsai M, Winter J, Emson PC, Woolf CJ (1994) Neurite outgrowth and GAP-43 mRNA expression in cultured adult rat dorsal root ganglion neurons: effects of NGF or prior peripheral axotomy. J Neurosci Res 39:634–645.

Inbal R, Rousso M, Ashur H, Wall PD, Devor M (1987) Collateral sprouting in skin and sensory recovery after nerve injury in man. Pain 28:141–154.

Isaacson LG, Crutcher KA (1998) Uninjured aged sympathetic neurons sprout in response to exogenous NGF in vivo. Neurobiol Aging 19:333–339.

Jackson PC, Diamond J (1981) Regenerating axons reclaim sensory targets from collateral nerve sprouts. Science 214:926–928.

Jackson PC, Diamond J (1983) Failure of intact cutaneous mechanosensory axons to sprout functional collaterals in skin of adult rabbits. Brain Res 273:277–283.

Jackson PC, Diamond J (1984) Temporal and spatial constraints on the collateral sprouting of low-threshold mechanosensory nerves in the skin of rats. J Comp Neurol 226:336–345.

Jenq CB, Jenq LL, Bear HM, Coggeshall RE (1988) Conditioning lesions of peripheral nerves change regenerated axon numbers. Brain Res 457:63–69.

Kalil RE, Schneider GE (1975) Abnormal synaptic connections of the optic tract in the thalamus after midbrain lesions in newborn hamsters. Brain Res 100:690–698.

Kalous A, Keast JR (2010) Conditioning lesions enhance growth state only in sensory neurons lacking calcitonin gene-related peptide and isolectin B4-binding. Neuroscience 166:107–121.

Ketschek A, Gallo G (2010) Nerve growth factor induces axonal filopodia through localized microdomains of phosphoinositide 3-kinase activity that drive the formation of cytoskeletal precursors to filopodia. J Neurosci 30:12185–12197.

Kinnman E (1987) Collateral sprouting of sensory axons in the hairy skin of the trunk: a morphological study in adult rats. Brain Res 414:385–389.

Kobayashi NR, Fan DP, Giehl KM, Bedard AM, Wiegand SJ, Tetzlaff W (1997) BDNF and NT-4/5 prevent atrophy of rat rubrospinal neurons after cervical axotomy, stimulate GAP-43 and Talpha1-tubulin mRNA expression, and promote axonal regeneration. J Neurosci 17:9583–9595.

Koltzenburg M, Wall PD, McMahon SB (1999) Does the right side know what the left is doing? Trends Neurosci 22:122–127.

Krimm RF, Davis BM, Albers KM (2000) Cutaneous overexpression of neurotrophin-3 (NT3) selectively restores sensory innervation in NT3 gene knockout mice. J Neurobiol. 43:40–49.

Lawson SN, Waddell PJ (1991) Soma neurofilament immunoreactivity is related to cell size and fibre conduction velocity in rat primary sensory neurons. J Physiol 435:41–63.

Lorenzana AO, Lee JK, Mui M, Chang A, Zheng B (2015) A Surviving Intact Branch Stabilizes Remaining Axon Architecture after Injury as Revealed by In Vivo Imaging in the Mouse Spinal Cord. Neuron 86, 947–954.

Lu P, Jones LL, Tuszynski MH (2005) BDNF-expressing marrow stromal cells support extensive axonal growth at sites of spinal cord injury. Exp Neurol 191:344–360.

Mannion RJ, Doubell TP, Gill H, Woolf CJ (1998) Deafferentation is insufficient to induce sprouting of A-fibre central terminals in the rat dorsal horn. J Comp Neurol 393:135–144.

Mahar, M., Cavalli, V., 2018. Intrinsic mechanisms of neuronal axon regeneration. Nat Rev Neurosci 19, 323–337.

McCarter GC, Levine JD (2006) Ionic basis of a mechanotransduction current in adult rat dorsal root ganglion neurons. Mol Pain 2:28.

McCarthy PW, Lawson SN (1997) Differing action potential shapes in rat dorsal root ganglion neurones related to their substance P and calcitonin gene-related peptide immunoreactivity. J Comp Neurol 388:541–549.

McLachlan EM, Janig W, Devor M, Michaelis M (1993) Peripheral nerve injury triggers noradrenergic sprouting within dorsal root ganglia. Nature 363:543–546.

McMahon SB, Armanini MP, Ling LH, Phillips HS (1994) Expression and coexpression of Trk receptors in subpopulations of adult primary sensory neurons projecting to identified peripheral targets. Neuron 12:1161–1171.

McMahon SB, Kett-White R (1991) Sprouting of peripherally regenerating primary sensory neurones in the adult central nervous system. J Comp Neurol 304:307–315.

McQuarrie IG (1985) Effect of conditioning lesion on axonal sprout formation at nodes of Ranvier. J Comp Neurol 231:239–249.

McQuarrie IG, Grafstein B, Gershon MD (1977) Axonal regeneration in the rat sciatic nerve: effect of a conditioning lesion and of dbcAMP. Brain Res 132:443–453.

Mearow KM, Kril Y, Diamond J (1993) Increased NGF mRNA expression in denervated rat skin. Neuroreport 4:351–354.

Michael GJ, Averill S, Nitkunan A, Rattray M, Bennett DL, Yan Q, Priestley JV (1997) Nerve growth factor treatment increases brain-derived neurotrophic factor selectively in TrkA-expressing dorsal root ganglion cells and in their central terminations within the spinal cord. J Neurosci 17:8476–8490.

Michael GJ, Averill S, Shortland PJ, Yan Q, Priestley JV (1999) Axotomy results in major changes in BDNF expression by dorsal root ganglion cells: BDNF expression in large trkB and trkC cells, in pericellular baskets, and in projections to deep dorsal horn and dorsal column nuclei. Eur J Neurosci 11:3539–3551.

Nixon BJ, Doucette R, Jackson PC, Diamond J (1984) Impulse activity evokes precocious sprouting of nociceptive nerves into denervated skin. Somatosens Res 2:97–126.

Patapoutian A, Reichardt LF (2001) Trk receptors: mediators of neurotrophin action. Curr Opin Neurobiol 11:272–280.

Pertens E, Urschel-Gysbers BA, Holmes M, Pal R, Foerster A, Kril Y, Diamond J (1999) Intraspinal and behavioral consequences of nerve growth factor-induced nociceptive sprouting and nerve growth factor-induced hyperalgesia compared in adult rats. J Comp Neurol 410:73–89.

Petruska, J.C., Barker, D.F., Garraway, S.M., Trainer, R., Fransen, J.W., Seidman, P.A., Soto, R.G., Mendell, L.M., Johnson, R.D., 2014. Organization of sensory input to the nociceptive-specific cutaneous trunk muscle reflex in rat, an effective experimental system for examining nociception and plasticity. J Comp Neurol 522, 1048–1071.

Phillips LL, Autilio-Gambetti L, Lasek RJ (1983) Bodians silver method reveals molecular variation in the evolution of neurofilament proteins. Brain Res 278:219–223.

Qiu J, Cafferty WBJ, McMahon SB and Thompson SWN (2005) Conditioning injury-induced spinal axon regeneration requires Signal Transducer and Activator of Transcription 3 activation. J. Neurosci. 25: 1645–1653.

Ramon y Cajal S (1928) Degeneration and regeneration of the Nervous System. London: Oxford University Press.

Richardson PM, Verge VM (1987) Axonal regeneration in dorsal spinal roots is accelerated by peripheral axonal transection. Brain Res 411:406–408.

Robinson PP (1988) Observations on the recovery of sensation following inferior alveolar nerve injuries. Br J Oral Maxillofac Surg 26:177–189.

Rosenzweig, E.S., Courtine, G., Jindrich, D.L., Brock, J.H., Ferguson, A.R., Strand, S.C., Nout, Y.S., Roy, R.R., Miller, D.M., Beattie, M.S., Havton, L.A., Bresnahan, J.C., Edgerton, V.R., Tuszynski, M.H., 2010. Extensive spontaneous plasticity of corticospinal projections after primate spinal cord injury. Nat Neurosci 13, 1505–1510.

Ruscheweyh R, Forsthuber L, Schoffnegger D, Sandkuhler J (2007) Modification of classical neurochemical markers in identified primary afferent neurons with Abeta-, Adelta-, and C-fibers after chronic constriction injury in mice. J Comp Neurol 502:325–336.

Schneider GE (1973) Early lesions of superior colliculus: factors affecting the formation of abnormal retinal projections. Brain Behav Evol 8:73–109.

Shi TJ, Tandrup T, Bergman E, Xu ZQ, Ulfhake B, Hokfelt T (2001) Effect of peripheral nerve injury on dorsal root ganglion neurons in the C57 BL/6J mouse: marked changes both in cell numbers and neuropeptide expression. Neuroscience 105:249–263.

Skene JH (1984) Growth-associated proteins and the curious dichotomies of nerve regeneration. Cell 37:697–700.

Smith DS, Skene JH (1997) A transcription-dependent switch controls competence of adult neurons for distinct modes of axon growth. J Neurosci 17:646–658.

Snider WD, Zhou FQ, Zhong J, Markus A (2002) Signaling the pathway to regeneration. Neuron 35:13–16.

Theriault E, Diamond J (1988) Nociceptive cutaneous stimuli evoke localized contractions in a skeletal muscle. J Neurophysiol 60:446–462.

Tobias CA, Shumsky JS, Shibata M, Tuszynski MH, Fischer I, Tessler A, Murray M (2003) Delayed grafting of BDNF and NT-3 producing fibroblasts into the injured spinal cord stimulates sprouting, partially rescues axotomized red nucleus neurons from loss and atrophy, and provides limited regeneration. Exp Neurol 184:97–113.

Tsujino, H., Kondo, E., Fukuoka, T., Dai, Y., Tokunaga, A., Miki, K., Yonenobu, K., Ochi, T., Noguchi, K., 2000. Activating transcription factor 3 (ATF3) induction by axotomy in sensory and motoneurons: A novel neuronal marker of nerve injury. Mol Cell Neurosci 15, 170–182.

Tucker BA, Rahimtula M, Mearow KM (2005) A procedure for selecting and culturing subpopulations of neurons from rat dorsal root ganglia using magnetic beads. Brain Res Brain Res Protoc 16:50–57.

Vavrek R, Girgis J, Tetzlaff W, Hiebert GW, Fouad K (2006) BDNF promotes connections of corticospinal neurons onto spared descending interneurons in spinal cord injured rats. Brain 129:1534–1545.

Wang X, Seed B (2003) A PCR primer bank for quantitative gene expression analysis. Nucleic Acids Res 31:e154.

Woolf CJ, Shortland P, Sivilotti LG (1994) Sensitization of high mechanothreshold superficial dorsal horn and flexor motor neurones following chemosensitive primary afferent activation. Pain 58:141–155.

Ygge J (1984) On the organization of the thoracic spinal ganglion and nerve in the rat. Exp Brain Res 55:395–401.

ZGraggen WJ, Fouad K, Raineteau O, Metz GA, Schwab ME, Kartje GL (2000) Compensatory sprouting and impulse rerouting after unilateral pyramidal tract lesion in neonatal rats. J Neurosci 20:6561–6569.

